# The NSL complex promotes neural development by preventing R-loop induced replication stress

**DOI:** 10.1101/2025.08.22.671281

**Authors:** Jolanthe Lingeman, Diana van den Heuvel, Jeroen van den Berg, Jenny K. Singh, Rosanne van Hooijdonk, Wouter W. Wiegant, Daphne E.C. Boer, Calvin S.Y. Lo, Andreas Panagopoulos, Román González-Prieto, Marnix Franken, Nikki Kolsters, Olivier G. de Jong, Alfred C.O. Vertegaal, Matthias Altmeyer, Nitika Taneja, Alexander van Oudenaarden, Martijn S. Luijsterburg, Nael Nadif Kasri, Brooke L. Latour, Haico van Attikum

**Affiliations:** Department of Human Genetics, Leiden University Medical Center, Leiden, The Netherlands; Oncode Institute, Hubrecht Institute-KNAW (Royal Netherlands Academy of Arts and Sciences) and University Medical Center Utrecht, Utrecht, The Netherlands; Department of Molecular Genetics, Erasmus MC Cancer Institute, Erasmus University Medical Center, Rotterdam, The Netherlands; Oncode Institute, Erasmus University Medical Center, Erasmus MC Cancer Institute, Rotterdam, the Netherlands; Department of Molecular Mechanisms of Disease, University of Zurich (UZH), Zurich, Switzerland; Department of Cell and Chemical Biology, Leiden University Medical Center, Leiden, The Netherlands; Department of Human Genetics, Radboud UMC, Donders Institute for Brain, Cognition and Behaviour, Nijmegen, The Netherlands; Department of Pharmaceutics, Utrecht University, Utrecht, The Netherlands

**Keywords:** Non-Specific Lethal (NSL) complex, DNA replication, R-loops, Neurodevelopment, Koolen-de Vries syndrome (KdVS)

## Abstract

Early neuronal development relies on the proliferation of neural progenitor cells, making this developmental stage particularly vulnerable to DNA replication impediments. Here, we identify a non-canonical role for the non-specific lethal (NSL) complex in safeguarding DNA replication during neurodevelopment. The NSL complex, which acetylates histone 4 at gene promoters and is mutated in Koolen-de Vries syndrome (KdVS), prevents unscheduled R-loop accumulation at weakly transcribed promoters that are typically devoid of R-loops. Single-cell sequencing reveals that loss of NSL causes replisome stalling and delayed S-phase progression. Neural organoids derived from KdVS patients exhibit impaired DNA replication, developmental abnormalities, and reduced synapse formation, driven entirely by unscheduled R-loop accumulation. These findings reveal that faithful DNA replication is critical for early neurodevelopment.

## Introduction

Human development and tissue maintenance require precise control of cell proliferation and differentiation. DNA replication, orchestrated by the cell cycle-regulated replisome, ensures accurate genome duplication and proper cell fate determination. This is especially critical for rapidly proliferating cells such as neural progenitor cells during early brain development^1–4^, which may explain why mutations in replisome components are increasingly linked to genetic disorders with neurodevelopmental abnormalities, including intellectual disability^5–9^. However, the nature of the replication defects and their causative link to neural dysfunction in such disorders remains elusive.

Koolen-de Vries syndrome (KdVS) is a multisystem, neurodevelopmental disorder that exhibits several phenotypic similarities with replication diseases, including intellectual disability, hypotonia, motor developmental delays, and congenital malformations^10, 11^. Other clinically relevant neurological features are epilepsy and structural brain abnormalities^12, 13^. KdVS results from a microdeletion of 17q21.31 that includes *KANSL1*, or from heterozygous mutations in *KANSL1*. The two genetic diagnoses are phenotypically indistinguishable, implicating *KANSL1* haploinsufficiency as the underlying cause of KdVS^14, 15^.

KANSL1 serves as the scaffold protein of the non-specific lethal (NSL) complex, a highly conserved chromatin remodeling complex comprising nine proteins^16, 17^. The core subunits of the NSL complex include KANSL1, KANSL2, KANSL3, and PHF20^17–19^. MOF, encoded by the *KAT8* gene, is a histone acetyltransferase that acetylates histone H4 at lysines 5 and 8, when associated with the NSL complex, thereby regulating transcription^17–24^. In neuronal models derived from individuals with KdVS, *KANSL1* haploinsufficiency leads to reduced NSL complex activity, resulting in transcriptional changes that disrupt autophagy and impair neuronal function by decreasing synaptic density^25^.

The NSL complex also has diverse nuclear functions beyond transcriptional regulation, including maintenance of nuclear architecture and genome integrity through acetylation of non-histone targets like Lamin A/C^18, 26^. Moreover, MOF has been implicated in promoting faithful DNA replication^27^. Similar to patients with KdVS, patients with MOF syndrome, which is caused by mutations in *KAT8*, display neural proliferation and differentiation abnormalities during neurodevelopment^28^. Although these findings suggest that loss of the NSL complex causes the observed neurodevelopmental abnormalities, it is unclear whether this results from the dysregulation of transcription, DNA replication, or both in neural cells.

Here we report a critical role for NSL in maintaining R-loop homeostasis and ensuring proper DNA replication during neurodevelopment. By utilizing KdVS patient-derived and engineered induced pluripotent stem cells (iPSCs) and applying single cell EdU-sequencing (scEdU-seq) and DNA-RNA immunoprecipitation followed by sequencing (DRIP-seq), we demonstrate that loss of the NSL complex leads to R-loop accumulation at lowly transcribed TSSs, leading to reduced DNA replication and prolonged S-phase progression. These insights provide a mechanistic link between defective R-loop regulation, disrupted DNA replication, and neurodevelopmental abnormalities.

## Results

### Haploinsufficiency of *KANSL1* reduces cell proliferation and DNA replication

The NSL complex is essential for human cell proliferation^19^, but the specific impact of *KANSL1* haploinsufficiency on the growth of cells remains unclear. To address this, we investigated the proliferation capacity of CRISPR/Cas9-engineerd heterozygous *KANSL1* (*KANSL1^+/-^)* and patient-derived induced pluripotent stem cells (iPSCs) from individuals with KdVS (Fig. 1A)^29^. Western blot analysis confirmed a reduction in KANSL1 protein levels in *KANSL1*-deficient iPSCs compared to controls (Extended Data Fig. 1A). To examine cell proliferation, we co-cultured mCherry-expressing *KANSL1^+/-^* or KdVS cells in a 1:1 ratio with enhanced green fluorescent protein (eGFP)-expressing control cells (Extended Data Fig. 1B). Subsequently, the ratio of mCherry to GFP was monitored over time using flow cytometry. Both *KANSL1^+/-^*and KdVS-derived iPSCs exhibited a significant reduction in proliferation when compared to controls (Extended Data Fig. 1C).

**Fig. 1.**
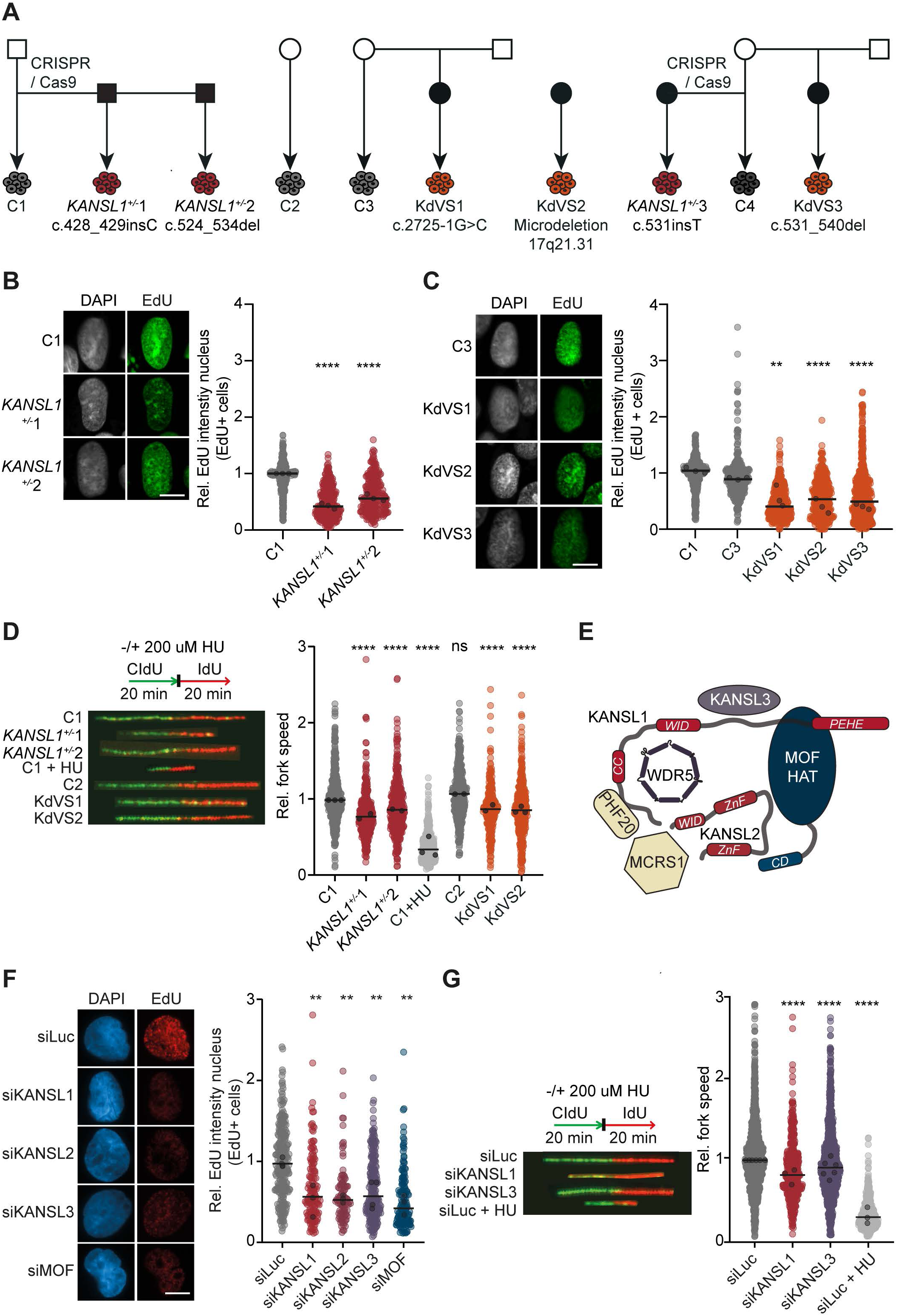
NSL loss reduces cell proliferation and DNA replication. A Pedigree of control and *KANSL1* haploinsufficient iPSC lines. Filled black squares and circles represent affected male and female lines, respectively. *KANSL1*^+/-^1 and *KANSL1^+/-^*2 CRISPR/Cas9-engineered iPSC lines are derived from C1, *KANSL1*^+/-^3 is derived from C4. Engineered lines carry heterozygous mutations in exon 2. C2 is an independent female control. C3 and C4 are maternal controls to KdVS1 and KdVS3, respectively. Patient lines KdVS1, KdVS2, and KdVS3 have heterozygous loss of *KANSL1*. Affecting variants are noted in the diagram. B Representative images (left) and relative EdU intensity (right) of EdU-positive *KANSL1*^+/-^ iPSCs. Median EdU levels were normalized to C1, which was set to 1. N=3 independent experiments. Black line represents the median of all data points, black circles the median of the independent experiments; one-way ANOVA with Dunnett’s multiple comparisons test on medians of independent experiments (**** p<0.0001). Scale bar 5 μm. C Representative images (left) and relative EdU intensity (right) of EdU-positive KdVS iPSCs. Median EdU levels were normalized to C1 and C3 combined, which was set to 1. N=3 independent experiments. Black line represents the median of all data points, black circles the median of the independent experiments; one-way ANOVA with Dunnett’s multiple comparisons test on medians of independent experiments (** p<0.01, **** p<0.0001). Scale bar 5 μm. D Representative images (left) and relative replication fork speed (right) from DNA fibers assays using *KANSL1* haploinsufficient and KdVS iPSC lines. Median fork speed was normalized to C1, which was set to 1. N=2-3 independent experiments. Black line represents the median of all data points, black circles the median of the independent experiments; Kruskall Wallis test with Dunn’s multiple comparisons test on all data points (ns=not significant, **** p<0.0001). E Schematic overview of the NSL complex^18^. F Representative images (left) and relative EdU intensity (right) of EdU-positive U2OS cells 6 days after treatment with the indicated siRNAs. Median EdU levels were normalized to siLuc, which was set to 1. N=3-4 independent experiments. Black line represents the median of all data points, black circles the median of the independent experiments; one-way ANOVA and Dunnett’s multiple comparisons test on median independent experiments (** p<0.01). Scale bar 10 μm. G Representative images (left) and relative replication fork speed (right) from DNA fiber assays with U2OS cells 4 days after treatment with the indicated siRNAs. N=3-6 independent experiments. Black line represents the median of all data points, black circles the median of the independent experiments; Kruskall Wallis test with Dunn’s multiple comparisons test on all data points (**** p<0.0001).

To investigate whether the reduced proliferation rates were due to impaired DNA replication, we pulsed *KANSL1^+/-^* and KdVS-derived iPSCs with the nucleotide analog 5-ethynyl-2′-deoxyuridine (EdU) and quantified the levels of EdU incorporation as a measure of DNA synthesis. Both *KANSL1^+/-^* and KdVS-derived iPSCs showed a significant reduction in EdU incorporation during S-phase compared to controls (Fig. 1B and 1C), indicating a decrease in DNA replication. Next, we performed DNA fiber analysis to explore whether this reduction was due to a decrease in replication fork speed. Indeed, we found that replication fork speed was lower in *KANSL1^+/−^* and KdVS-derived iPSCs relative to controls (Fig. 1D). These findings suggest that *KANSL1* haploinsufficiency compromises DNA replication, contributing to the reduced proliferation observed for KdVS patient cells.

### Loss of the NSL complex impairs cell proliferation and DNA replication

Loss of KANSL1, a scaffold protein of the NSL complex (Fig. 1E), impairs NSL’s function in transcription regulation^16, 18, 25^. We therefore sought to determine whether the reduced proliferation and DNA replication capacity of *KANSL1* haploinsufficient cells could be attributed to loss of NSL function. To this end, we used siRNA-mediated knockdown to individually deplete three core components of the NSL complex, KANSL1, KANSL2 and KANSL3, as well as its catalytic subunit MOF, in human osteosarcoma U2OS cells. Depletion of these NSL components, which was confirmed by western blot analysis and reverse transcriptase quantitative PCR (RT-qPCR), reduced cell proliferation compared to cells treated with a control siRNA (Extended Data Fig. 1D and 1E). These results are in accordance with previous work implicating the NSL complex in cell cycle regulation^19, 30^. Next, we assessed DNA replication and found that depletion of KANSL1, KANSL2, KANSL3 or MOF reduced EdU incorporation by approximately 50% or more (Fig. 1F). Furthermore, similar to KdVS patient-derived iPSCs, KANSL1- and KANSL3-depleted U2OS exhibited a moderate, but significant decrease in replication fork speed, as measured by DNA fiber analysis (Fig. 1G). Together, these results demonstrate that loss of the NSL complex impairs DNA synthesis in both U2OS cells and KdVS patient-derived iPSCs, contributing to reduced cell proliferation.

### DNA replication in early replicating regions is impaired by loss of the NSL complex

DNA replication dynamics, including origin firing and replication fork speed, are precisely coordinated to assure high fidelity duplication of the genome^31, 32^. To further assess how DNA replication is affected by loss of the NSL complex, we examined DNA replication dynamics by performing single cell EdU-seq (scEdU-seq) on U2OS control cells, and cells depleted of KANSL1 or KANSL3 (Fig. 2A and Extended Data Fig. 2A-D)^33^. While global DNA replication timing remained largely unchanged in KANSL-depleted cells compared to control cells (Fig. 2A), a closer examination of DNA replication tracks from early S-phase, specifically the first 25% of S-phase, revealed a lower density of scEdU-seq signal in KANSL-depleted cells than in controls cells. In addition, we found that these specific regions are replicated much later in S-phase (Fig. 2A). Remarkably, in KANSL-depleted cells, the genome-wide DNA replication timing during mid-S phase closely resembled the early replication patterns observed in control cells. (Extended Data Fig. 2E). This suggests that DNA replication in early replication domains takes much longer in KANSL-depleted cells, consistent with the reduced replication speeds observed in DNA fiber assays (Fig. 1G).

**Fig. 2.**
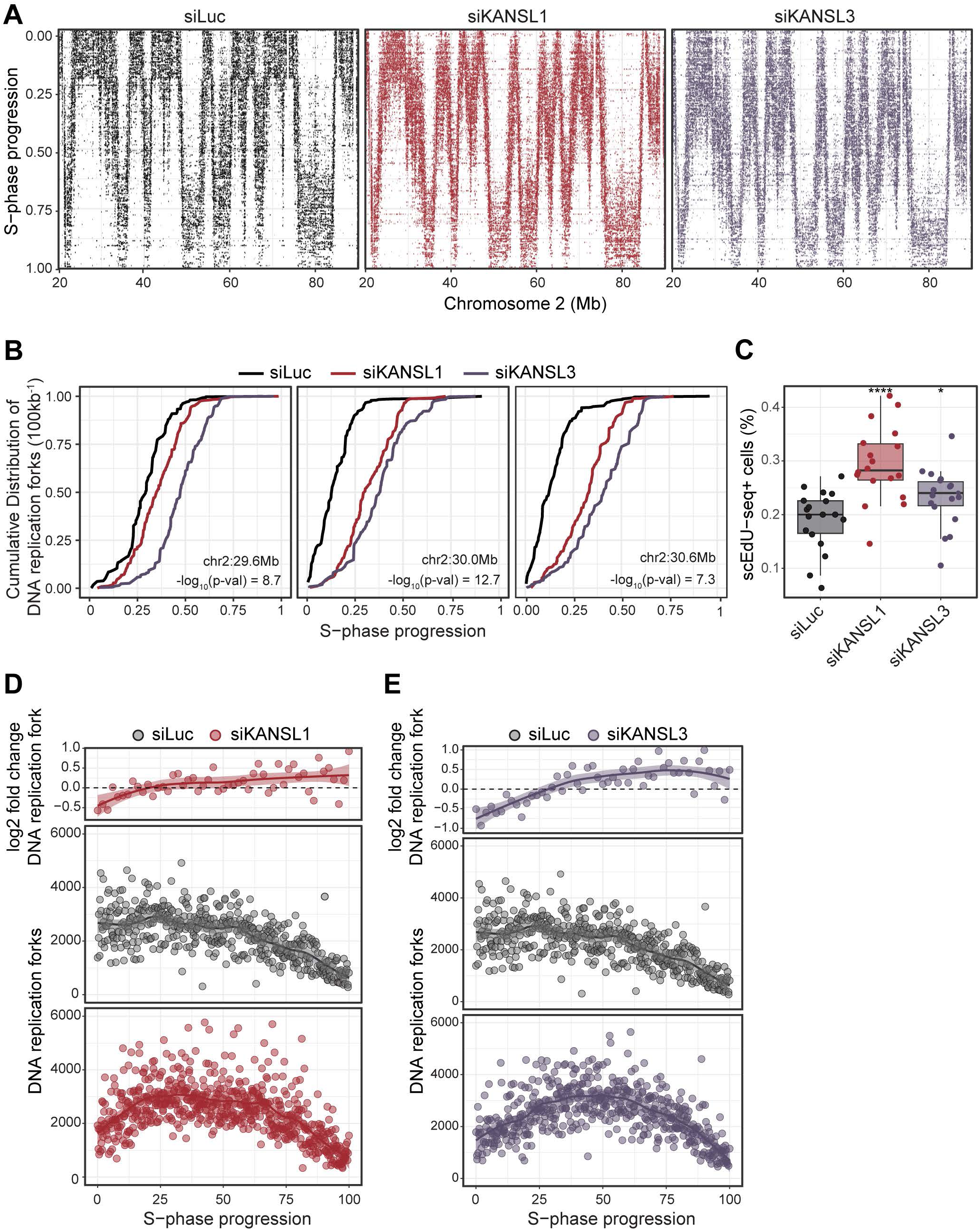
scEdU-seq reveals that early replicating regions are affected by NSL loss. A Heatmap of DNA replication forks from U2OS cells 4 days after treatment with the indicated siRNAs ordered according to S-phase progression (y-axis) and position in chromosome 2 (x-axis) for 70 megabases of chromosome 2 assayed with scEdU-seq. B Cumulative frequency distribution (x-axis) of DNA replication forks over S-phase progression in cells from A. Genomic location and -log10(adj.p-val) are labeled in each plot. C The percentage of scEdU-seq positive cells from A with each dot indicating a technical replicate (i.e. 384-well plate). The box of the boxplot is defined by the median ± IQR and whiskers are 1.5X IQR. Two-tailed t-test with multiple testing correction was performed (* p<0.05, **** p<0.0001). D The number of DNA replication forks (y-axis) for individual single siLuc- and siKANSL1-treated U2OS cells from A plotted over S-phase progression (x-axis) (bottom plots), and the log2 fold change of DNA replication forks of siKANSL1/siLuc (*y* axis) plotted over S-phase progression (*x* axis, top plot). E As in D, except for siKANSL3-treated U2OS cells.

To quantify the increase in DNA replication time per genomic region, we computed a cumulative frequency distribution of the number of forks during S-phase progression (Fig. 2B) and assessed their effect size and significance (Extended Data Fig. 2F and 2G). As expected, we identified several regions where DNA replication is initiated at the start of S-phase progression. However, the replication of these regions in KANSL-depleted cells lagged behind that of control cells. This suggests that in the absence of KANSL1 or KANSL3, these regions are harder to replicate, resulting in delayed replication, a prolonged S-phase, and the completion of DNA replication occurring later in the S-phase^34^. Consistent with a prolonged S-phase, we observed an increase in the percentage of EdU-positive KANSL1- and KANSL3-depleted cells (Fig. 2C).

Next, we asked if there is an adaptive response to the lag in DNA replication of early replicating regions. To assess this, we quantified the number of active replication forks. We found that KANSL-depleted cells display a lower number of forks in early S-phase (Fig. 2D, 2E and Extended Data Fig. 2H), which is in line with the decreased density of replication tracks (Fig. 2A). However, the number of forks increased in mid to late S-phase in these cells, likely to compensate for the initial delay in DNA replication in early S-phase (Fig. 2D, 2E and Extended Data Fig. 2H). Taken together, our results demonstrate that loss of the NSL complex impairs replication of early-replicating regions. To compensate, NSL-depleted cells increase their fork numbers in mid-S phase, offsetting the reduced number of active forks in early S-phase.

### Replication fork stalling increases in the absence of the NSL complex

To determine whether the impaired DNA replication in early S-phase KANSL-depleted cells is caused by obstacles that block fork progression, we examined the symmetry of bidirectional forks using DNA fiber analysis. Strikingly, we found that depletion of KANSL1 or KANSL3 reduced fork symmetry, resembling the reduction observed after treatment with a low dose of hydroxyurea, which perturbs DNA replication by depletion of dNTP pools^35^ (Fig. 3A and 3B). Moreover, a marked reduction in fork symmetry was also observed in *KANSL1^+/-^* and KdVS patient-derived iPSCs (Fig. 3C). To further assess replication fork stalling, we examined the impact of KANSL3 depletion on replication fork stalling following hydroxyurea treatment of cells. KANSL3 loss led to an increase in stalled replication forks, accompanied by heightened origin firing (Fig. 3D). This phenotype was similar to that observed in RAD51-depleted cells, which was included as a positive control^36, 37^.

**Fig. 3.**
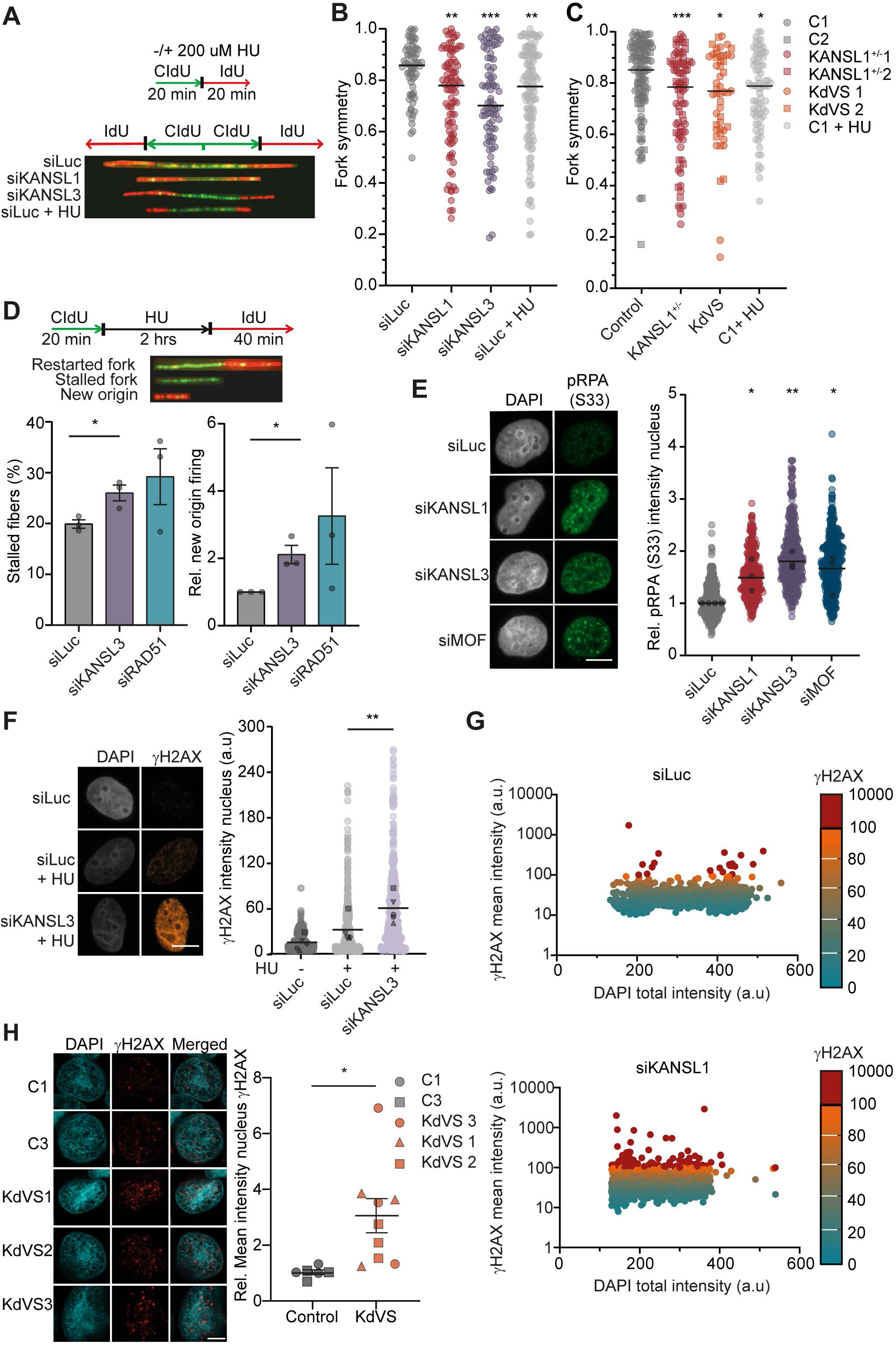
Replication forks stall in absence of the NSL complex. A Labelling scheme and representative images of DNA fiber assay assessing replication fork symmetry in KANSL1- and KANSL3-depleted U2OS cells. Cells treated with 200 mM HU during CldU and IdU labelling are a positive control. B Representative images (left) and fork symmetry analysis (right) in U2OS cells 4 days after treatment with the indicated siRNAs. N=3 independent experiments. Black line represents the median of all data points; Kruskall Wallis test with Dunn’s multiple comparisons test on all data points (**p<0.01, *** p<0.001). C Fork symmetry analysis in control, *KANSL1^+/-^*and KdVS iPSCs. Data of multiple cell lines grouped per category. N=2-3 independent experiments per cell line. Black line represents the median of all data points; Kruskall Wallis test with Dunn’s multiple comparisons test on all data points (*p<0.05, *** p<0.001). D Labelling scheme (top) and representative images of DNA fiber assay assessing replication fork stalling and new origin firing in HU-treated U2OS cells (bottom). Fork stalling and relative new origin firing in U2OS cells exposed to HU 2 days after treatment with the indicated siRNAs. Mean levels of new origin firing were normalized to siLuc, which was set to 1.N=3 independent experiments. Mean ± SEM shown, black circles represent means of independent experiments; unpaired two-tailed t-test (* p<0.05)). E Representative images (left) and relative nuclear pRPA intensity(right), phosphorylated at serine 33, in U2OS cells, 2 days after treatment with the indicated siRNAs. pRPA levels were normalized to siLuc, which was set to 1. N=3-4 independent experiments. Black line represents the median of all data points, black circles the median of the independent experiments; one-way ANOVA and Dunnett’s multiple comparisons test on median independent experiments (* p<0.05, ** p<0.01). Scale bar 10 μm. F Representative images (left) and γH2AX levels (right) in U2OS cells, treated for 4 hours +/-2mM HU, 2 days after treatment with the indicated siRNAs. N=5 independent experiments. Black line represents the mean of all data points, black circles the mean of the independent experiments; paired t-test on means independent experiments between siLuc and siKANSL3 treated with HU (* p<0.05, ** p<0.01). Scale bars 10 μm. G γH2AX and DAPI intensity measured by high-throughput immunofluorescence in U2OS cells, 6 days after treatment with the indicated siRNAs. A representative experiment of N=2 independent experiments is shown. H Representative nuclei (left) and γH2AX mean fluorescence intensity per nucleus (right) across control and KdVS iPSCs. Two control cell lines and three patient cell lines were analysed and grouped per category. KdVS data were normalized to C1 and C3, which was set to 1. N=3 independent experiments per cell line. Each symbol represents a mean of an independent experiment; mean ± SEM; unpaired two-tailed t-test (* p<0.05). Scale bar 5 µm.

Stalled replication forks can undergo fork reversal, which generates an end that is amenable to nucleolytic degradation. In line with previous work, we found that BRCA2 protects reversed replication forks against degradation^38^. However, KANSL3-depletion did not impact fork degradation (Extended Data Fig. 3A), suggesting that KANSL-deficient cells primarily suffer from replication fork stalling rather than from the reversal and degradation of stalled forks.

Supporting this notion, we found that KANSL-depleted cells were hypersensitive to replication fork stalling induced by hydroxyurea treatment (Extended Data Fig. 3B-E). This phenotype was rescued by ectopic expression of siRNA-resistant GFP-KANSL3, indicating it is not an off-target effect of the siRNA that was used to deplete KANSL3 (Extended Data Fig. 3D and 3E). Moreover, ATR kinase is activated in response to stalled replication forks, where it phosphorylates multiple target proteins, including replication protein A (RPA) and histone H2AX^39^. In line with increased replication fork stalling, we observed increased levels of ATR-phosphorylated RPA (S33) and H2AX (γH2AX) in KANSL- and MOF-depleted cells (Fig. 3E-G and Extended Data Fig. 3F-G), as well as in KdVS patient-derived iPSCs (Fig. 3H). Collectively, these findings indicate that DNA replication is hampered by replication fork stalling in KANSL-depleted U2OS cells and KdVS patient-derived iPSCs.

### Loss of the NSL complex causes R-loops to accumulate

To identify the origin of replication fork blockages observed in KANSL-depleted cells, we performed mass spectrometry analysis. This analysis showed that KANSL3 interacts strongly with other NSL complex members, but not with DNA replication factors, suggesting it may influence DNA replication indirectly (Extended Data Fig. 4A). The NSL complex regulates transcription, a process that contributes to the formation of three-stranded nucleic acid structures called R-loops^18, 19, 23, 40^. R-loops form when an RNA strand hybridizes with one strand of DNA, displacing the other strand as a single-stranded loop. While they play important roles in gene regulation, R-loops can also cause genomic instability by obstructing the replisome and inducing replication fork stalling^41–43^. We therefore assessed DNA-RNA hybrid formation by immunofluorescence using the S9.6 antibody, which preferentially binds DNA-RNA hybrids in R-loops. Strikingly, we detected an increase in S9.6 signal intensity in KANSL1- and MOF-depleted U2OS cells (Fig. 4A and Extended Data Fig. 4B), as well as in KdVS patient-derived iPSCs (Fig. 4B). Importantly, the increased S9.6 signal in KANSL3-depleted U2OS cells was rescued by overexpression of siRNA-resistant GFP-KANSL3 (Fig. 4C and 4D), indicating it is not an off-target effect of siRNA. Moreover, increased R-loop levels were also observed in KANSL3-depleted retinal pigment epithelial (RPE-1) cells, indicating it is not a cell type-specific effect (Extended Data Fig. 4C and 4D).

**Fig. 4.**
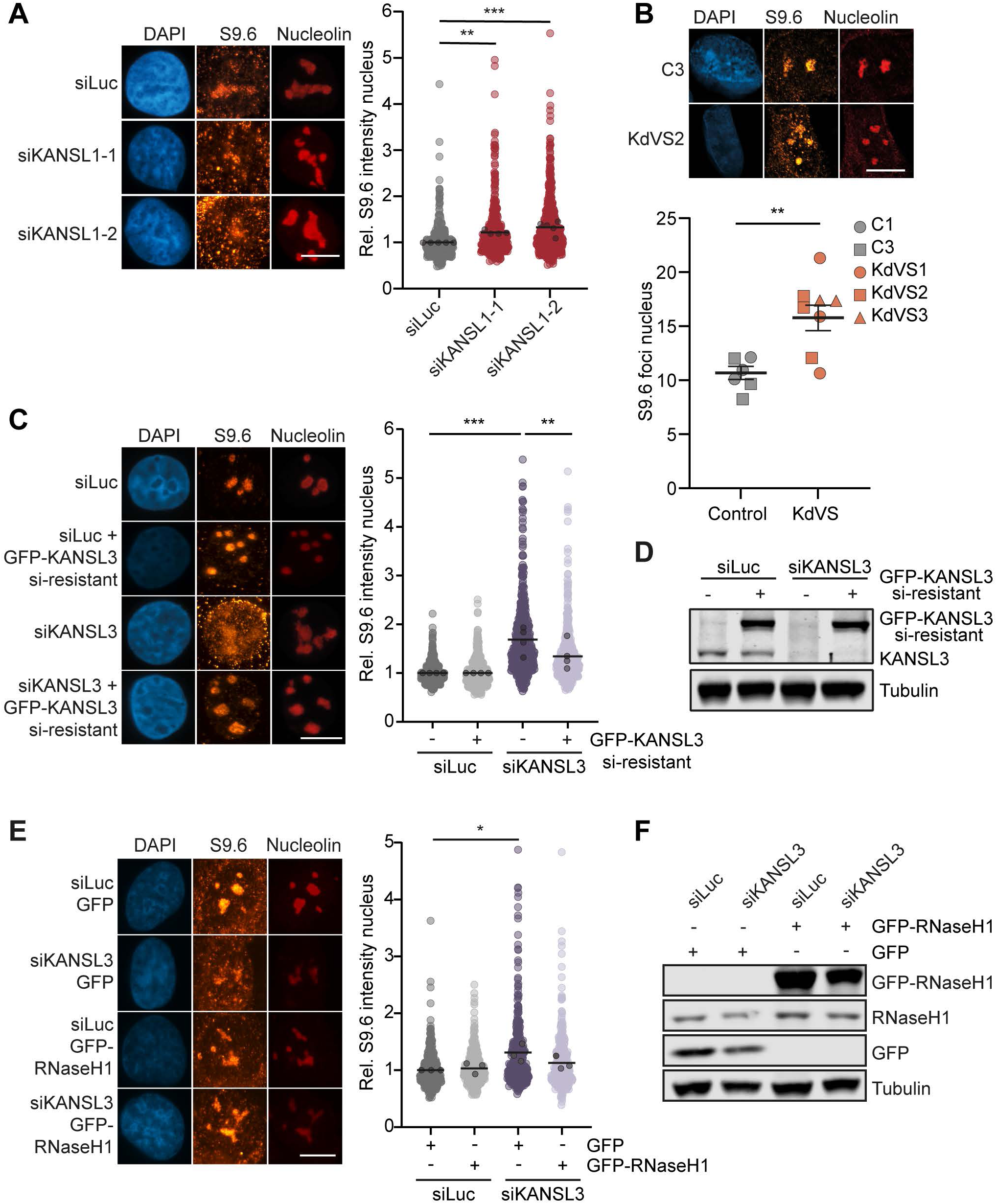
R-loop accumulation in KdVS and NSL depleted cells. A Representative nuclei (left) and R-loop analysis (Right) in U2OS cells, 2 days after treatment with the indicated siRNAs. R-loops are imaged by the S9.6 antibody and analysed as mean intensity per nucleus excluding the nucleolar signal. S9.6 levels were normalized to siLuc, which was set to 1. N=4-5 independent experiments. Black line represents the mean of all data points, black circles the means of the independent experiments; one-way ANOVA and Dunnett’s multiple comparisons test on means independent experiments (** p<0.01, *** p<0.001). Scale bar 10 μm. B Representative nuclei (left) and R-loop analysis (right) of control and KdVS patient iPSCs, imaged by the S9.6 antibody and analysed as the number of S9.6 foci per nucleus. Two control cell lines and three patient cell lines were analysed and grouped per category. N=3 independent experiments per cell line. Symbols represent the mean of a cell line in an experiment; mean ± SEM; unpaired two-tailed t test (** p< 0.01). Scale bar 10 μm. C Representative nuclei (left) and R-loop analysis (right) in U2OS cells 4 days after treatment with the indicated siRNAs and -/+ siRNA resistant GFP-KANSL3. R-loops are imaged by the S9.6 antibody and mean intensity per nucleus, excluding nucleolar signal, is analysed. S9.6 levels were normalized to siLuc -/+ siRNA resistant GFP-KANSL3, which was set to 1. N=4 independent experiments. Black line represents the mean of all data points, black circles the means of the independent experiments; paired two-way ANOVA (** p<0.01, *** p<0.001). Scale bar 10 μm. D Western blot indicating KANSL3 protein expression levels of recue experiments described in C. Endogenous KANSL3 and overexpression of GFP-KANSL3 siRNA resistant shown. Tubulin is shown as loading control. One representative experiment of two experiments shown. E Representative nuclei (left) and rescue of R-loop phenotype (right) in U2OS cells, 4 days after treatment with the indicated siRNAs and lentiviral transduction with GFP or GFP-RNaseH1. S9.6 levels were normalized to siLuc GFP, which was set to 1. N=3 independent experiments. Black line represents the mean of all data points, black circles the means of the independent experiments; paired two-way ANOVA (* p<0.05). Scale bar 10 μm. F Western blot indicating GFP, GFP-RNaseH1 and endogenous RNaseH1 protein expression levels of rescue experiments described in E. Tubulin is shown as loading control. One representative experiment of two experiments shown.

Although the S9.6 antibody detects R-loops, it also binds dsRNA and ssRNA, albeit with lower affinity^44^. We therefore verified its specificity in our assays. First, overexpression GFP-tagged RNaseH1, an enzyme that degrades the RNA strand in R-loops^45^, reduced the increase in S9.6 signal in KANSL3-depleted cells, whereas GFP alone had no effect (Fig. 4E and 4F). Second, incubation of cells with mutated recombinant GFP-RNaseH1 protein (GFP-RNaseH1-D210N), which is able to bind but not degrade R-loops^44^, showed an increase in nuclear GFP-RNaseH1-D210N signals following depletion of KANSL3 or Senataxin (SETX), which was used as a positive control due to its known role as a helicase in R-loop processing (Extended Data Fig. 4E)^42, 44, 46^. Finally, consistent with the notion that R-loops can arise from replication stress, we observed particularly high R-loop levels following hydroxyurea treatment of KANSL3-depleted cells (Extended Data Fig. 4F)^47^. Taken together, these results show that the NSL complex suppresses the formation of R-loops that could potentially obstruct DNA replication.

### R-loops accumulate at weak promoters in the absence of NSL

To determine where R-loops accumulate in the absence of KANSL3, we performed genome-wide DRIP-seq. U2OS cells were transfected in triplicate with either siLuc or siKANSL3, followed by immunoprecipitation of R-loops using the S9.6 antibody. The immunoprecipitated material was treated or not treated with recombinant RNaseH1, sonicated, prepared for library construction, and sequenced. Peak calling revealed that R-loops primarily mapped to protein-coding regions, especially gene promoters (transcription start sites; TSS), 5’ UTRs and terminators (transcription termination sites; TTS) (Fig. 5A), as previously described^48^. Importantly, R-loop signals were completely abolished by treatment with RNaseH1.

**Fig. 5.**
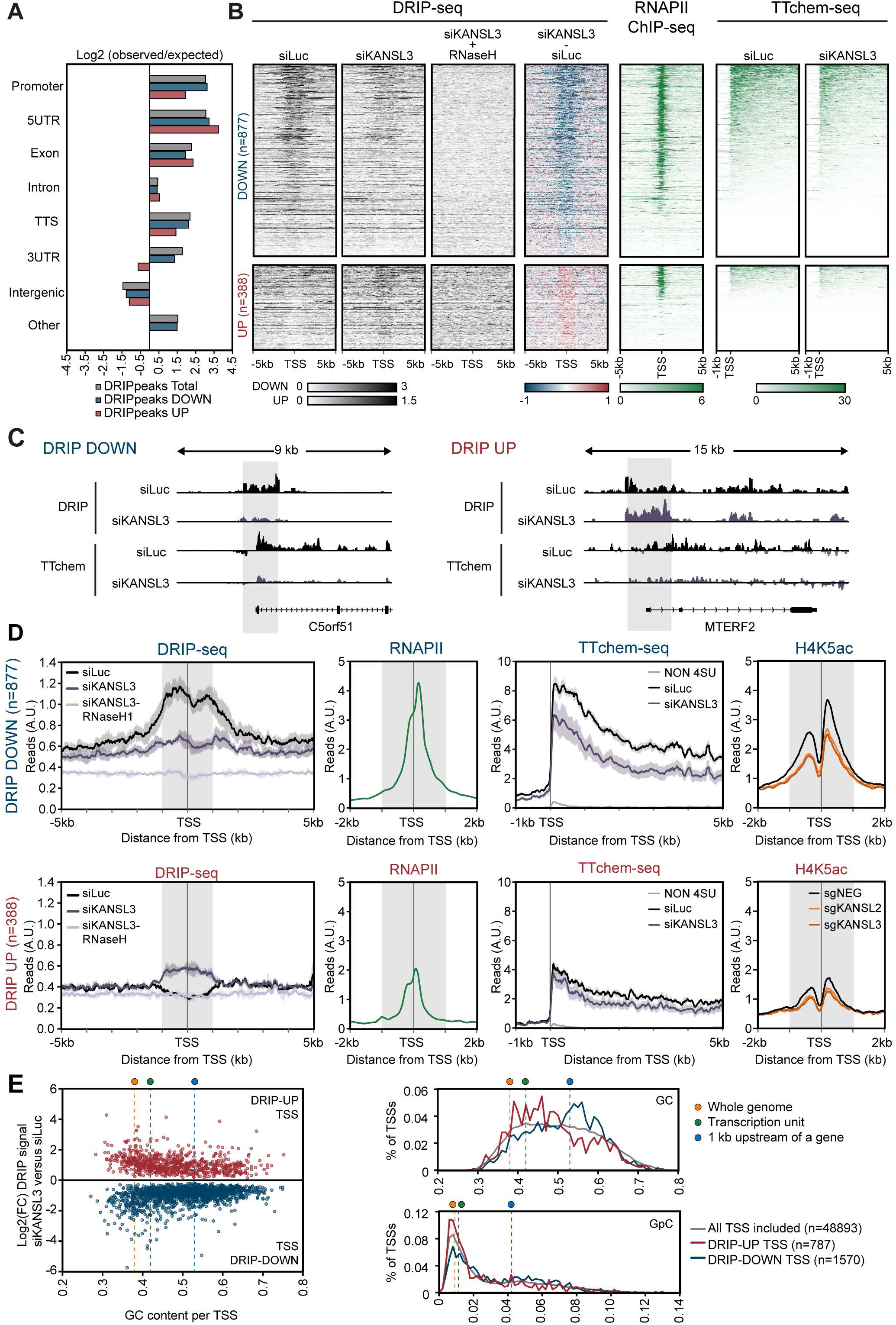
R-loops accumulate at weak promoters in absence of NSL. A Distribution of identified DRIP-peaks within different genomic regions in U2OS cells, 4 days after treatment with the indicated siRNAs. Presented are all identified DRIP peaks (N=20129), DRIP-peaks with decreased (DOWN N=1899) or increased (UP N=179) signal upon KANSL3 depletion compared to siLuc. B Heatmaps of DRIP-seq, RNAPII ChIP-seq and TTchem-seq signals at indicated regions around TSSs of genes of at least 5kb that show decreased (DRIP-DOWN) or increased (DRIP-UP) DRIP signal at their TSS (+/-1 kb) upon KANSL3 depletion compared to controls. C Genome track examples of DRIP-seq and TTchem-seq signals at a TSS with decreased (DOWN) or increased (UP) DRIP-seq signal in siKANSL3 versus siLuc. TTchem-seq signals were separated in sense (positive track) and anti-sense (negative track) signals. Grey shaded areas are identified DRIP-peak regions. D Metaprofiles of DRIP-seq, RNAPII ChIP-seq, TTchem-seq and H4K5ac ChIP-seq signals around TSSs with decreased (DOWN) or increased (UP) DRIP signal in siKANSL3 compared to siLuc. The grey shaded areas are regions from -1kb to +1kb around the TSS used to quantify decreased or increased DRIP signal. E GC and CpG content of TSSs (-/+ 1kb) that show decreased (blue) or increased (red) DRIP signal in siKANSL3 versus siLuc. Data is presented as individual TSSs relative to the Log2 fold change of DRIP signal in siKANSL3 versus siLuc (left) or as histogram representation (right). Average GC or CpG contents for the whole genome, transcription units or 1kb upstream of a gene were obtained for comparison^76^.

Since the NSL complex promotes transcription initiation at housekeeping genes, partly through the acetylation of H4K5 and H4K8^19^, we examined whether KANSL3 knockdown affects R-loops at gene promoters. Reduced transcription initiation would typically be expected to decrease co-transcriptional R-loops^49^. However, our imaging experiments revealed an increase in nuclear R-loop levels. Following KANSL3 depletion, we observed a decrease in R-loops at 1,570 promoters, while an increased signal was detected at 759 promoters. To prevent quantification bias by short genes, we focused on promoters of genes at least 5 kb in length (n=877 DOWN and n=388 UP) (Fig. 5B). A typical gene track of a DOWN and UP promoter is shown (Fig. 5C). Metaprofiles of all DOWN or UP promoters demonstrated consistent changes across all three replicates, with signals fully abolished by RNaseH1 treatment (Fig. 5D).

Next, we generated metaprofiles for the DOWN and UP genes using our previously published RNAPII ChIP-seq data ^50^, and published datasets for H4K8ac and H4K5ac (Fig. 5D, Extended Data Fig. 5A, and 5B)^19^. In addition, we performed RNA-seq, using a transient transcriptome sequencing protocol in combination with chemical fragmentation of the nascent RNA (TTchem-seq)^51^, in U2OS cells transfected with either siLuc or siKANSL3 (Fig. 5D, Extended Data Fig. 5C, and 5D). Strikingly, in control cells, the 877 genes with reduced R-loop levels upon KANSL depletion (DOWN genes) exhibited high RNAPII binding and nascent transcription at their promoters, along with high levels of H4K8ac and H4K5ac (Fig. 5D and Extended Data Fig. 5B). These results suggest that the DOWN genes are highly transcribed genes. Following KANSL3 depletion, a marked reduction in nascent RNA levels, as well as in H4K5 and H4K8 acetylation was observed for these genes (Fig. 5D, Extended Data Fig. 5B, 5E, and 5F). This suggests that the DOWN genes are NSL target genes that lose co-transcriptional R-loops due to reduced transcriptional output.

In contrast to the DOWN genes, the 388 genes with increased R-loop levels upon KANSL depletion (UP genes) displayed low RNAPII binding and at least two-fold lower nascent transcription levels in control cells, with correspondingly low levels of H4K8ac and H4K5ac, all of which were only minimally affected by KANSL3 depletion (Fig 5D and S5B). Despite this, these genes specifically accumulated R-loops at their promoters. TTchem-seq analysis confirmed that the increase in R-loops for these 388 UP genes was not due to enhanced transcription in either the sense or antisense direction (Extended Data Fig. 5C and 5G). Strikingly, however, we noted a higher GC content in the UP genes compared to the whole genome, which is a defining feature of R-loop-prone genomic regions^52^, though not as pronounced as in the R-loop regions of the DOWN genes (Fig. 5E). We propose that these R-loops are more persistent and, therefore, more easily detected using our imaging methods. Possibly due to the low transcriptional flux through these regions. This is conceptually similar to the global loss of nascent transcription and co-transcriptional R-loops detected by DRIP-seq after treatment with the splicing inhibitor Pladienolide B^53^, which at the same time triggers an increase in persistent R-loops detectable by imaging methods^54–56^.

### Loss of the NSL complex induces R-loop-dependent fork stalling and impairs DNA replication

Our DRIP-seq revealed elevated R-loop levels around TSSs in KANSL-depleted cells, while scEdU-seq indicated delayed initiation of DNA replication in early-replicating regions during S-phase. (Fig. 2B, e.g. 29-30Mb). Based on these findings, we investigated whether R-loops contribute to the delayed replication of early-replicating regions. To this end, we integrated the median replication timing for 100 kilobase bins in the first 20% of S-phase, as measured by scEdU-seq, with the significant DRIP-seq UP and DOWN signals (Fig. 5B). We found that in KANSL-depleted cells, the timing of replication in regions that gained R-loops at TSSs is delayed, while this was not seen genome-wide in regions with elevated R-loops. In addition, replication timing in KANSL-depleted cells was neither affected in regions that lost R-loops at TSSs, nor in regions with reduced R-loop levels genome-wide (Fig. 6A and 6B). These results suggest that R-loops hinder replication timing in KANSL-depleted cells, potentially by impeding DNA replication through replication fork stalling^5, 41, 42^. To test this, we addressed whether the increased R-loop levels contribute to fork stalling in KANSL3- and MOF-depleted cells. *In situ* proximity ligation assays (PLA) assay were performed using the S9.6 antibody to label R-loops and EdU incorporation to detect newly synthesized DNA. An increase in PLA foci was observed in KANSL3- and MOF-depleted cells, indicating a marked stalling of replication forks at persistent R-loops (Fig. 6C). Based on this, we examined whether overexpression of RNaseH1 in KANSL-depleted cells would prevent fork stalling and overcome the observed DNA replication defects. Indeed, GFP-RNaseH1 overexpression not only rescued the fork symmetry defects caused by KANSL1 and KANSL3 loss (Fig. 6D), but also restored overall DNA synthesis, as measured by EdU incorporation in replicating cells (Fig. 6E). Collectively, our findings suggest that the NSL complex prevents R-loop-dependent replication fork stalling at weak promoters, thereby ensuring timely DNA replication of these genomic regions.

**Fig. 6.**
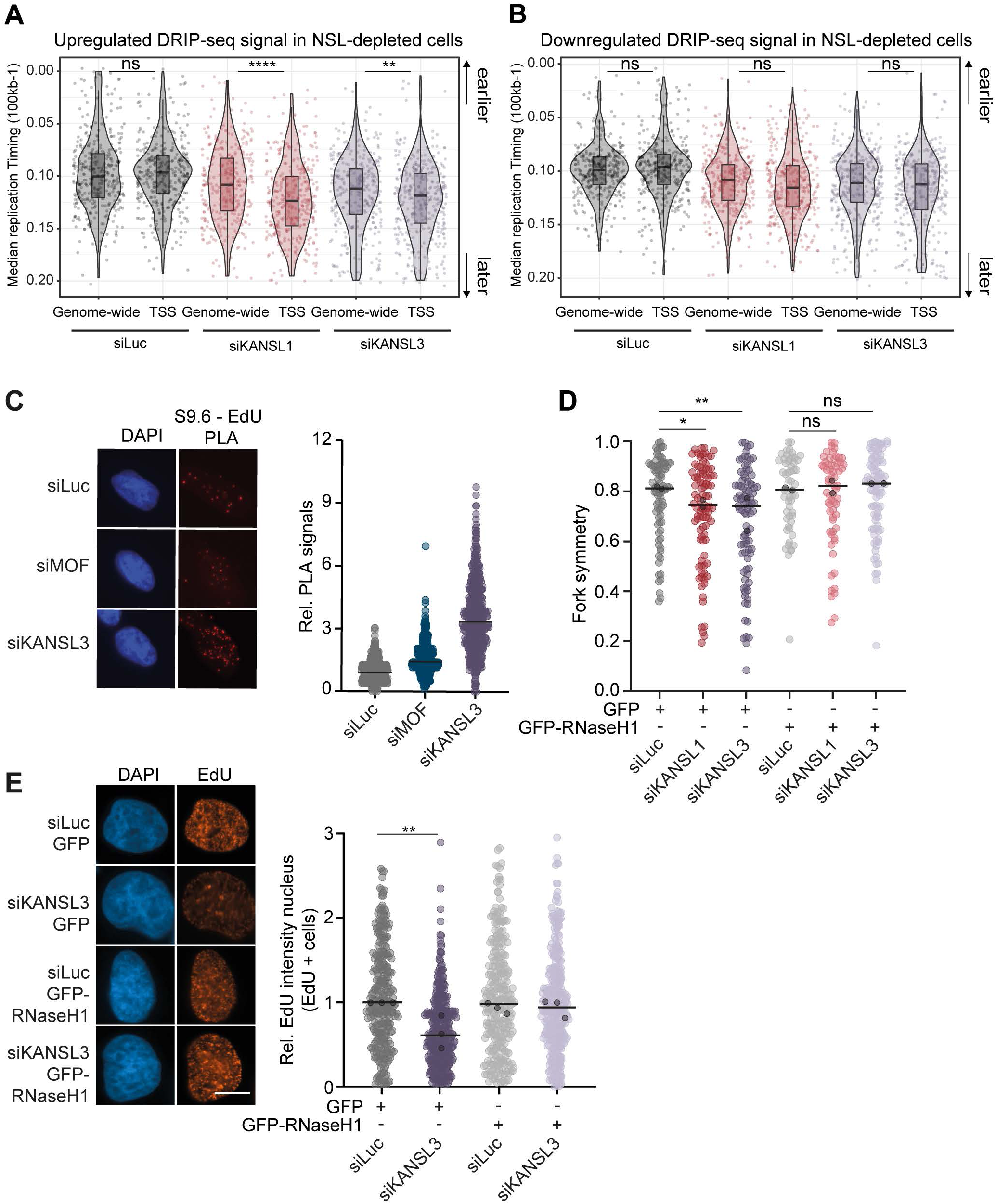
R-loops cause replication fork stalling. A Median DNA replication timing (y-axis) for 100kb bins in U2OS cells, 4 days after treatment with the indicated siRNAs, overlapping with indicated regions derived from upregulated peaks from DRIP-seq (x-axis). The box of the boxplot is defined by the median ± IQR and whiskers are 1.5X IQR; Wilcoxon rank sum test was performed (ns= not significant, ** p<0.01, **** p<0.0001). B Median DNA replication timing (y-axis) for 100kb bins in U2OS cells, 4 days after treatment with the indicated siRNAs, overlapping with indicated regions derived from downregulated peaks from DRIP-seq (x-axis). The box of the boxplot is defined by the median ± IQR and whiskers are 1.5X IQR; Wilcoxon rank sum test was performed (ns=not significant). C Representative nuclei (left) and quantification of proximity ligase assay (PLA) (right) examining collisions between the replication fork and DNA-RNA hybrids in MRC-5 human fibroblasts 2 days after treatment with the indicated siRNAs. Newly synthesized DNA is visualized by EdU incorporation, DNA-RNA hybrids are imaged with the S9.6 antibody. PLA levels were normalised to siLuc, which was set to 1. Data of one representative experiment of two experiments shown. Black line represents the median of an experiment; one experiment of two representative experiments shown. D Rescue of replication fork symmetry phenotype measured by DNA fiber assay in U2OS cells, 4 days after treatment with the indicated siRNAs and lentiviral transduction of GFP or GFP-RNAseH1. N=2 independent experiments. Black line represents the median of all data points, black circles the medians of the independent experiments; Kruskall Wallis test on all data points (ns= not significant, *p<0.05, ** p<0.01). E Representative nuclei (left) and rescue of DNA synthesis (right) measured by EdU incorporation in EdU positive cells in 4 days after treatment with the indicated siRNAs and lentiviral transduction of GFP or GFP-RNAseH1. EdU intensity was normalized to siLuc GFP, which was set to 1. N=3 independent experiments. Black line represents the median of all data points, black circles the medians of the independent experiments; unpaired two-way ANOVA on the individual medians (** p<0.01). Scale bar 10 μm.

### *KANSL1* haploinsufficiency results in DNA replication defects and affects neuronal development

Neurodevelopmental abnormalities, including structural brain malformations such as enlarged ventricles, heterotopia, and agenesis or dysgenesis of the corpus callosum, are a hallmark of KdVS^13^. However, it remains unclear whether R-loop induced DNA replication defects contribute to these malformations. To investigate this, we differentiated *KANSL1*^+/-^ and KdVS-derived patient iPSCs into neural organoids (Extended Data Fig. 6A). These neural organoids developed specific neural architecture, including structures resembling the ventricular zone (VZ), subventricular zone (SVZ), and cortical plate (CP), along with specific PAX6+ apical and HOPX+ outer radial glia neural progenitor cell types, as well as differentiating TBR1+ and CTIP2+ neurons (Extended Data Fig. 6B). Moreover, *KANSL1* haploinsufficient neural organoids demonstrated typical macroscopic neural development, with no issues in organoid formation or morphology when compared to controls (Extended Data Fig. 6C). To evaluate whether the replication-stress phenotype observed in *KANSL1* haploinsufficient iPSCs contributes to KdVS-associated neurodevelopmental abnormalities, we examined DNA replication in organoid-derived neural progenitor cells during their self-renewal phase. To this end, we isolated SOX2+ radial glia cells from 16-days old neural organoids, coinciding with the onset of neurogenesis in neural organoids^57^, and assessed EdU incorporation in this population using flow cytometry (Extended Data Fig. 6D). *KANSL1* haploinsufficient radial glia cells exhibited reduced EdU incorporation (Fig. 7A and 7D), indicating a reduction in DNA replication in *KANSL1* haploinsufficient neural progenitor cells.

**Fig. 7.**
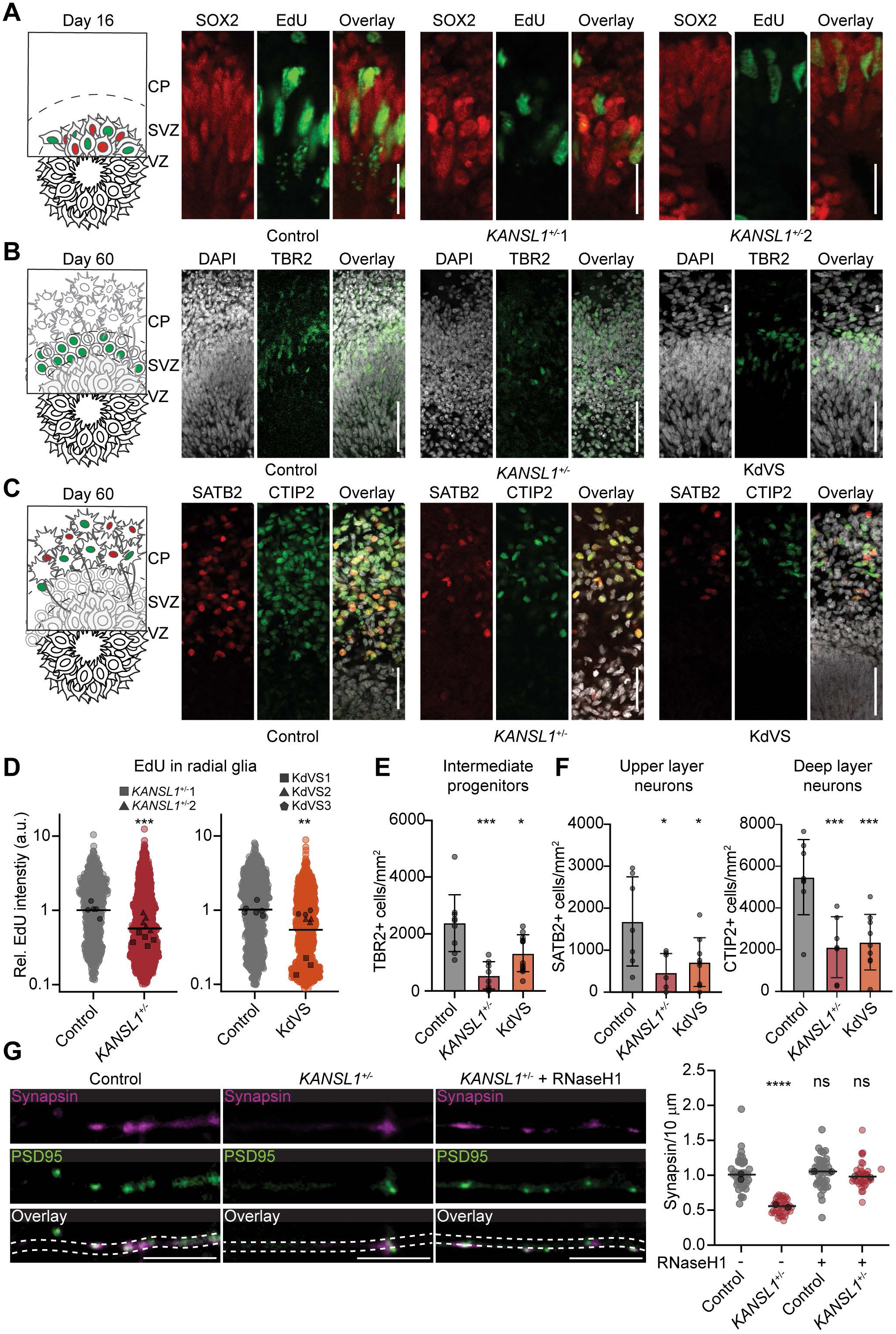
Neurodevelopment is disrupted in *KANSL1* haploinsufficient organoids and iNeurons. A Graphical insert (left) and representative images (right) of neuroepithelial rosettes in C1, *KANSL1*^+/-^1, and *KANSL1*^+/-^2 day 16 organoids stained for the radial glia marker SOX2 (red) and EdU (green). Graphical insert depicts the red SOX positive radial glia nuclei within the ventricular zone (VZ) with sparse green EdU labeling. Scale bar 25 μm. B Graphical insert (left) and representative images (right) of DIV60 organoids stained for the intermediate progenitor marker TBR2 (in green) localized to the VZ and subventricular zone (SVZ). Graphical insert depicts the green TBR2+ intermediate progenitor cells in the SVZ. Scale bars 50 μm. C Graphical insert (left) and representative images (right) of DIV60 organoids stained for the deep layer excitatory neuron marker CTIP2 (green) and the upper layer excitatory neuronal marker SATB2 (red). Graphical insert depicts the red SATB2+ and green CTIP2+ excitatory neurons in the cortical plate (CP). Scale bars 100 μm. D Relative EdU incorporation in control and *KANSL1*^+/-^ (left) and control and KdVS (right) organoids. Organoids were treated with EdU for 4 hours and then dissociated for single cell flow cytometry analysis of EdU incorporation. EdU integration was assessed in radial glia cells by gating on the live, SOX2+ population. EdU levels were normalized to control cells, and which was set to 1. N=3-5 organoids. Black line represents the median of all cells, black shapes the medians of individual organoids; One-way ANOVA and Dunnett’s multiple comparisons test for significance was performed on medians per organoid (** p<0.01, *** p<0.001). E Quantification of intermediate progenitors via TRB2+ cells in C4, *KANSL1*^+/-^3, and KdVS 3 organoids. Each black circle represents a cross section. N=5 organoids. Mean ±SD shown; Kruskal-Wallis with multiple comparisons test for significance was performed on all organoid sections (* p<0.05, ***p<0.001). F Quantification of deep layer CTIP2+ (left) and superficial layer SATB2+ (right) excitatory neurons in C4, *KANSL1*^+/-^3, and KdVS 3 organoids. Each black circle represents a cross section. N=5 organoids. Mean ±SD shown; one-way ANOVA and Dunnett’s multiple comparisons test for significance was performed on all organoid sections (* p<0.05, *** p<0.001). G Representative images (left) and number of active excitatory synapses per 10 μm of primary dendrite (right) of primary dendrites from day 21 iNeurons. iNeurons were cultured for 21 days and left untreated or transduced with RNaseH1 lentivirus on day 14. iNeurons were stained for the presynaptic marker Synapsin (magenta) and the post synaptic marker PSD95 (green), and the dendritic marker MAP2 (white). A functional synapse is defined as colocalization between synapsin and PSD95. N=2 independent experiments. Black line represents the median of all data points, black circles the medians of the independent experiments; Kruskal-Wallis test with Dunn’s multiple comparisons test (ns=not significant, **** p<0.0001). Scale bars 10 μm.

To explore the potential downstream effects of dysregulated replication dynamics, we next examined differentiation trajectories in *KANSL1* haploinsufficient neural organoids. In day 60 organoids, we observed a reduction in intermediate (TBR2+) progenitor cells (Fig. 7B and 7E). As TBR2+ intermediate progenitors primarily give rise to excitatory pyramidal neurons, this prompted us to assess the impact on pyramidal neuron formation^58^. Strikingly, both early-born deep-layer (CTIP2+) and later-born superficial-layer (SATB2+) neurons were reduced in *KANSL1⁺/⁻* and KdVS organoids (Fig. 7C and 7F). These findings align with recent studies linking dysregulated R-loop levels to defects in excitatory neuron development^59^.

Given the specific depletion of excitatory pyramidal neurons, we next evaluated whether *KANSL1* haploinsufficiency affected excitatory neuron maturation by assessing synapse formation in iPSC-derived neurons (iNeurons) (Extended Data Fig. 7A). By day 21 of differentiation, *KANSL1*^+/-^ neurons exhibited a significant reduction in functional synapse number, consistent with previous observations in KdVS patients and *KANSL1* haploinsufficient iNeurons^25^ (Fig. 7G). Notably, overexpression of RNaseH1 rescued the synaptic defect when introduced on day 14 of differentiation. Importantly, RNaseH1 overexpression had no substantial effect on synapse number in control (C1) iNeurons (Fig. 7G and Extended Data Fig. 7B-D). These results indicate that R-loops delay synaptic maturation in *KANSL1* haploinsufficient neurons. Collectively, our findings suggest that disrupted R-loop homeostasis leads to DNA replication defects and contributes to the neurodevelopmental deficits observed in KdVS (Extended Data Fig 7E).

## Discussion

While DNA replication deficits are implicated in human disease, the exact mechanisms linking them to disease etiology, particularly neurodevelopmental defects, remain largely unknown. KdVS is a rare disorder caused by *KANSL1* haploinsufficiency that shares several features with replication-associated diseases, including neurodevelopmental defects^6, 13–15^. The work described here shows that KANSL1/NSL prevents the accumulation of R-loops at TSSs with low basal transcription and H4K5/H4K8 acetylation, where R-loops are typically absent. In the absence of a functional NSL complex, these R-loops delay DNA replication in early S-phase by stalling replisomes, leading to aberrant DNA replication initiation, prolonged S-phase progression, and reduced cell proliferation. As a result, neural organoids derived from KdVS patients, which have reduced KANSL1 levels and NSL activity, exhibit disrupted DNA replication alongside defects in neurodevelopmental differentiation, while *KANSL1* haploinsufficient neurons display reduced synapse formation in an R-loop-driven manner. These findings provide new insights into the pathobiology of KdVS, revealing that R-loop-induced DNA replication stress contributes to the neurodevelopmental defect in KdVS, suggesting it is a replication disease (Extended Data Fig. 7E).

### R-loop homeostasis and DNA replication control are intimately linked

R-loops are more likely to form in regions with high transcriptional activity. When transcription and replication occur simultaneously at the same loci, R-loops can directly hinder progression of the replication machinery through head-on or co-transcriptional collisions^49^. However, in KANSL-depleted cells, delayed replication was mainly found at weak promoters with low transcription and acetylation activity (Fig. 5D), yet increased R-loop levels, suggesting this phenomenon is not related to NSL’s transcriptional function or its histone acetyltransferase activity. Indeed, recent work showed that R-loops, but not transcription machineries, are the cause of transcription-replication collisions, particularly in the head-on orientation^60^. Moreover, replisomes can modulate R-loop levels based on orientation, suppressing R-loops in the co-directional orientation, while enhancing their formation in the head-on orientation. Our scEdU-seq experiments, however, revealed not only an R-loop-dependent delay in early replication, but also a reduced number of replication forks at the start of S-phase in KANSL-depleted cells, indicative of deregulated origin firing (Fig. 2D and 2E). Deregulated origin firing and subsequent replication stress exacerbate head-on collisions, further promoting R-loop accumulation. Supporting this notion, we observed increased R-loop levels, pRPA and γH2AX in KANSL3-depleted cells experiencing HU-induced replication stress (Fig. 3F, Extended Data Fig. 3F, and 4G). Although, DNA replication initiation is hampered in KANSL3-depleted cells, replisomes that do fire reach TSSs prone to R-loop formation, thereby enhancing the levels of these structures. These persistent R-loops detected by the immunofluorescence and DRIP-seq analysis, likely form physical barriers that stall replisome, further delaying DNA replication (Extended Data Fig. 7E). In response to this replication stress, the ATR/CHK1 pathway is activated, particularly following head-on collisions, to halt the cell cycle and resolve stalled replication forks^60^. Consistently, we observed elevated levels of ATR-phosphorylated RPA (S33) and gH2AX (Fig. 3E-G and Extended Data Fig. 3E-F), along with increased replication fork stalling (Fig. 3D), suggesting that the ATR/CHK1 pathway may also indirectly contribute to replisome stalling and DNA damage formation in KANSL3-depeleted cells (Extended Data Fig. 7E). While these findings provide new insights into NSL’s role in controlling the intimate link between R-loop homeostasis and DNA replication, several questions remain. It is unclear, for instance, how the NSL complex controls DNA replication initiation and prevents the formation of persistent R-loops at specific TSSs.

### R-loops as regulators of transcription and DNA replication in the developing brain

Reduced synaptic transmission has been previously described in KdVS patient derived iNeurons and in the 17q21.31 microdeletion mouse model of KdVS^25, 61^. Our findings not only confirm these observations, but also demonstrate that the defect in synapse maturation is likely due to aberrant R-loop formation (Fig. 7G). Transcription can influence R-loop levels, especially around the TSS^62–64^. Conversely, R-loops can regulate transcription, playing a role in both gene activation and silencing^43, 65–70^. CpG islands and gene promotors favor R-loop formation, as R-loops protect against DNA methylation, prevent the formation of heterochromatin, and stimulate transcription^66–68^. Furthermore, lncRNA form R-loops with regulatory functions on transcription, both to stimulate and repress transcription^43, 62^. Recent work revealed an important role of R-loops in regulating neuronal gene expression^59^. These results were confirmed, by showing that neural progenitor cell-specific R-loops in TSSs arise mostly in GC skewed, early replicating and transcriptionally active genes, which are enriched for neural development^71^. Our DRIP-seq analysis revealed a GC skew for persistent R-loops at TSSs that impact DNA replication in U2OS cells (Fig. 5A). Similarly, such R-loops may delay DNA replication at TSSs in neural cells, impacting their maturation and/or function. In line with this, we observed increased levels of DNA damage following DNA replication perturbation in NSL-depleted U2OS cells and KdVS patient-derived iPSCs. A correlation between R-loops and DNA breaks in neural progenitor cells has been reported^71, 72^, but requires further investigation to resolve the mechanistic link between R-loop-induced DNA replication impediments, DNA damage induction and neurodevelopment.

### KdVS: R-loops, transcription and DNA replication disease

KdVS results from haploinsufficiency of *KANSL1*, impacting NSL-dependent histone H4 acetylation (H4K5/K8) and transcription regulation. Consequently, the developmental abnormalities observed in KdVS are closely linked to dysregulation of gene expression, particularly of genes crucial for neurodevelopment ^25, 61^. Here, we present evidence of DNA replication defects in NSL-depleted cells, including KdVS patient-derived cells, caused by alterations in R-loop homeostasis. R-loop-induced DNA replication impediments were observed at weak promoters with low transcription and H4K5/8 acetylation, suggesting they arise largely independent of transcription. Furthermore, these effects may underlie the neurodevelopmental abnormalities observed in the brain of KdVS patients. These findings align with recent studies linking DNA replication to neurodevelopmental disorders, such as Seckel syndrome and Meier-Gorlin syndrome^6, 9, 73–75^. Similar to KdVS patients, patients suffering from *KAT8* haploinsufficiency also display developmental and neurological impairments^28^. Given that MOF regulates transcription by H4 acetylation, likely including that of neuronal genes, and has been implicated in promoting DNA replication and cell proliferation^27, 28^, it is tempting to speculate that its loss also affects neurodevelopment by disrupting both transcriptional regulation and DNA replication. Taken together, this suggests that the neurodevelopmental defects in KdVS may be caused by a combination of transcription dysfunction and DNA replication defects. Characterizing the interplay between these processes in patients with KdVS, MOF syndrome, and other replication-related diseases with neurodevelopmental manifestations, will enhance our understanding of their roles in disease etiology.

## Supporting information

Lingeman_et_al_Supplementary_data

## Declaration of generative AI and AI-assisted technologies in the writing process

During the preparation of this work the author(s) used ChatGPT / OpenAI in order to improve language and readability. After using this tool/service, the author(s) reviewed and edited the content as needed and take(s) full responsibility for the content of the publication.

## Data and code availability

Both raw and processed NGS data were deposited in the Gene Expression Omnibus (GEO) under GEO: GSE289456 (reviewer token ifkpsugarvkthyj; scEdu-seq), GSE290178 (shczygqynfwhhyj; DRIP-seq), and GSE290179 (reviewer token ivubeamihpqndgx; TTchem-seq). Both raw and processed mass spectrometry data were deposited in the Protein Identification Database (PRIDE) under PXD061077 (reviewer token Iuw4OWaweGAC). All data will be publicly available as of the date of publication. Microscopy or western blot data reported in this paper can be shared by the lead contact upon request. This study did not involve the creation of original code. The software used for data analysis is detailed in the respective methods sections of the supplementary information file. Any additional information needed to reanalyze the data presented in this paper is available upon request from the lead contact.

## Methods

### Cell culture

iPSCs were cultured on Matrigel (Corning, 356237) or on human laminin (Biolamina, L521) in Essential 8 (E8) flex supplemented (Life Technologies, A2858501) with primocin (0.1 µg/ml; Invivogen, ANT-PM-05). Ngn2+ iPSCs were cultured on Matrigel (Corning, 356237) in E8 flex supplemented (Life Technologies, A2858501) with primocin (0.1 µg/ml; Invivogen, ANT-PM-05), puromycin (0.5 µg/ml) and G418 (0.5 µg/ml; Sigma-Aldrich, 4727878001). All iPSC lines were cultured at 37°C, 5% CO2. Medium was refreshed every 2 days and cells were passaged every 4-5 days, using an enzyme-free reagent (ReLeSR; Stem Cell Technologies, 05872). C1 is UCSFi001-A/wtc11 from the Coriell Institute for Medical Research and C2 is HPSI0314i-hoik_1 purchased from the HipSci human iPSC initiative. Generation of the C3, C4, KdVS1, KdVS2, KdVS3, and *KANSL1*^+/-^3 iPSC lines has previously been published^25^. C1, C3, KdVS1, KdVS2, and KdVS3 were determined to be karyotypically normal via high resolution karyotyping KaryoStat+ Karyotyping Service (ThermoFisher). C4 and *KANSL1*^+/-^3 were found to have acquired the commonly observed 20q11.21 gain. *KANSL1*^+/-^1 and *KANSL1*^+/-^2 were genomically assess by ddPCR (StemGenomics) and standard karyotyping (SCTC, Radboudumc) after CRISPR/Cas9 genomic editing, and Ngn2+ C1 and *KANSL1*^+/-^1 were genomically assessed with ddPCR (StemGenomics) after rtTA/Ngn2 stabilization. All karyotypically normal cell lines were used within 10 passages of genomic integrity assessment.

Human U2OS, U2OS GFP, U2OS dsRED, U2OS GFP-KANSL3 siRNA resistant, RPE1-hTERT, HEK293T, HCT116 and HCT116 GFP-KANSL3 siRNA resistant cells were cultured in 5% CO2 at 37°C in DMEM (Dulbecco’s modified Eagle’s medium) supplemented with 10% fetal calf serum and antibiotics. MRC-5 human fibroblasts were cultured in a 1:1 ratio of Dulbecco’s modified Eagle’s medium (DMEM) and Ham’s F10 (Invitrogen) supplemented with 10% fetal calf serum (FCS, Biowest) and 1% Penicillin/Streptavidin (PS, Sigma-Aldrich). All cell lines were regularly tested for mycoplasma infection and consistently found mycoplasma-free.

Generation of HCT116 Flp-In/T-Rex and U2OS Flp-In/T-Rex cells was described previously^77^. HCT116 Flp-In/T-Rex and U2OS Flp-In/T-Rex cells, which were generated using the Flp-In/T-Rex system (ThermoFisher Scientific), were a gift of Bradley Wouters (Princess Margaret Cancer Centre, Canada) and Geert Kops (University Medical Center Utrecht, the Netherlands). These cells were used to stably express inducible versions of GFP-NLS, as well as siRNA-resistant GFP-KANSL3WT by co-transfection of pCDNA5/FRT/TO-Puro plasmid encoding GFP or GFP-tagged KANSL3 (5 μg), together with pOG44 plasmid encoding the Flp recombinase (1 μg). After selection on 1 μg/mL puromycin, single clones were isolated and expanded. Both HCT116 Flp-In/T-Rex clones and U2OS Flp-In/T-Rex were incubated with 2 μg/mL doxycycline for 24-48h to induce expression of cDNAs.

### Transfections, siRNAs and plasmids

Cells were transfected with siRNAs using RNAiMAX (Invitrogen) according to the manufacturer’s instructions. Cells were transfected once with siRNAs 24 hours after seeding at a concentration of 40 nM and analyzed 48-72 hours after transfection unless otherwise indicated. siRNA sequences are listed in Supplementary Table 1.

Cells were transfected with plasmid DNA using Lipofectamine 2000 (Invitrogen) according to the manufacturer’s instructions and analyzed 24-48 hours after transfection.

The expression vector for full length human KANSL3 (64775: from Addgene; originally from Joan Conaway and Ronald Conaway), was amplified and cloned into pCDNA5/FRT/TO-Puro as a HindIII/KpnI fragment.siKANSL3-resistant KANSL3 cDNA was generated by introducing the underlined mutations CGACGATAACCTTAGGATCAG by overlap PCR and cloned as HindIII/KpnI fragment into pCDNA5/FRT/TO-Puro-KANSL3-WT. All KANSL3 expression constructs were verified using Sanger sequencing. GFP and GFP-RNaseH1 lentiviral overexpression constructs were a gift from the Luijsterburg lab (Leiden University Medical Center, the Netherlands).

### Genomic editing of iPSCs

Genome editing and characterization of C1 was performed by the Radboudumc Stem Cell Technology Center. In short, iPSCs were dissociated into single cells suspension and electroporated (4D nucleofector System X unit) with RNP complexes to establish a CRISPR/Cas9 edited cell pool, 48 hours after nucleofection single cells were seeded on a 10 cm dish coated with Matrigel (Corning, 356237). The remaining cells are used for sanger sequencing in order to determine the editing efficiency of the pool using the ICE software (Synthego), TIDER and the IDT analysis program. Seven days after seeding, the 48 clones (24 per sgRNA) were selected and screened via sanger sequencing to determine if the desired edit was present. The successfully edited clones that were expanded and characterized via karyotyping and assessed for trilineage differentiation and pluripotency with immunofluorescence microscopy. Specific targeting sequences for gRNAs are provided in Supplementary Table 1.

### Organoid differentiation

iPSCs were differentiated into neural organoids using the STEMdiff Cerebral Organoid kit (Stemcell Technologies, 08570) according to the manufacturer protocol with minor modifications. Specifically on day 0, 2, and 4 EB formation medium containing STEMdiff Cerebral Organoid Basal Medium 1 and STEMdiff Cerebral Organoid Supplement A was supplemented with 10 µM Y-27632 (Selleckchem, S1049), 2.5 µM dorsomorphin (Tocris, 3092) and 10 µM SB431542 (Stemcell Technologies, 72234).

### iNeuron differentiation

iPSCs were generated and differentiated into excitatory cortical iNeurons by doxycycline induced overexpression of *Neurog2* as previously described^78^, with minor modifications. For rescue experiments cells were treated once with 2 uL of concentrated RNase H1 lentivirus on either day 7 or day 14.

### Western blotting

iPSCs were plated at a density of 350.000 (control cells) and 500.000 (KANSL1 haploinsufficent cells) in a 6 well-plate coated with human laminin 521 or matrigel. The iPSC were harvested 48h after plating in RIPA buffer (pH 7.5) containing 150 mM NaCI (Sigma-Aldrich), 50 mM Tris-HCI (Invitrogen), 1% NP-40, 0.1% SDS, 0.5% Sodium Deoxycholate, 1mM EDTA (Sigma-Aldrich), and supplemented with protease inhibitor HALT (Fisher, #10127963). Protein concentrations were determined with the use of PierceTM BCA protein assay (Life Technologies, #23225). For each sample 10 ug was loaded and separated on 4-15% Mini-PROTEAN TGX Stain-Free gels (Bio-Rad, #4568084). The proteins were transferred to nitrocellulose membrane (Bio-Rad, 1704158). For blocking, the membranes were incubated with 5% non-fat dry milk (Santa Cruz Biotechnology, #sc-2325) in 0.1% PBS-T for 1h at room temperature. Primary antibodies that were used are KANSL1 (1:500; Sigma-Aldrich, HPA006874) and beta-actin (1:5000; Invitrogen, MA1-140). The antibodies were diluted in 5% non-fat dry milk and incubated overnight at 4°C. For visualization, the blots were incubated horseradish peroxidase-conjugated secondary antibody (HRP) for 1h at room temperature: goat anti-mouse (1:50000; Jackons ImmunoResearch, 115-035-062) and goat anti-rabbit (1:50000; Invitrogen, G21234). After incubation, blots were washed and incubated for 5 minutes at room temperate with ECL reagent (SuperSignalTM West Femto Maximum Sensitiviy Substrate, ThermoFisher Scientific, #34095) and imaged using ChemiDocTM Touch Gel Imaging System (Bio-Rad). The protein bands were quantified using ImageJ Software. Beta-actin was used as loading control.

Western Blotting in U2OS cells was performed as described previously^79^. Cells were lysed in 2x Laemmli buffer and boiled for 5-10 min at 95°C. Proteins were separated by SDS-PAGE using 4-12% pre-cast polyacrylamide gels (Criterion TGX pre-cast midi protein gel; Biorad or cat# NW04125; Invitrogen) and MOPS running buffer (cat#B000102; Invitrogen). Next, proteins were transferred onto nitrocellulose membranes (cat#AAWP02500; Millipore). Membranes were blocked in blocking buffer (Rockland or 5% milk solution in PBS-T (1x PBS with 0.1% Tween-20)). Protein expression was analyzed by immunoblotting with the indicated primary antibodies in blocking buffer (Rockland or 2% milk solution in PBS-T) (Supplementary Table 2) and secondary CF680 goat anti-rabbit or CF770 goat anti-mouse Ig antibodies (1:10000, Biotium). Membranes were scanned and analyzed using a Licor Odyssey scanner (LI-COR Biosciences).

### Competition assay

Stable lines expressing mCherry or eGFP were generated from all iPSC lines. To control for potential grow advantages introduced by lentiviral transduction, cells lines were assessed as both eGFP control versus mCherry *KANSL1* haploinsufficent line and mCherry control versus eGFP *KANSL1* haploinsufficient line. Flurorescent positive (FP+ Control and FP+ *KANSL1* haploinsufficient cells were plated in a 1:1 ratio on day 0 and assessed on day 3, 5, and 7 via flow cytometry (Beckman Coulter Gallios - 10 color) (Supplementary Fig. 1A). Proliferation capacity was assessed via the ratio between mCherry / eGFP positive cells. *KANSL1^+/-^1 and KANSL1^+/-^2* were assessed with C1 and KdVS1, 2, and 3 were assessed with C3.

U2OS dsRED cells, transfected with siLuc, were mixed in a 1:1 ratio with U2OS GFP cells transfected with siLuc, siKANSL1, siKANSL2, siKANSL3 and siMOF. The cells were mixed and counted 24h after siRNA transfection (day 1) and cultured for 3, 6 or 9 days. After culturing, dsRED and GFP cells were counted via Novocyte flow cytometer (Agilent) (Supplementary Fig. 1B). Proliferation capacity was determined as the ratio between GFP/dsRED positive cells. This was calculated based on cell counts of the GFP and dsRED positive cells and normalized to Day 1 and siLuc.

### DNA synthesis via EdU incorporation

iPSCs were plated as single cells at 150 cells/mm^2^ in µ-plate 96 well square glass bottom plates (ibidi GmbH, 89627). 40 hours after plating cells were pulsed with 20 uM of EdU for 4 hours, washed once with DPBS and then fixed. Wells were washed three times with PBS and then incubated in blocking buffer (PBS, 5% normal goat serum [Invitrogen, 10000 C], 5% normal horse serum [Gibco, 26050088], 5% normal donkey serum [Jackson Immuno Research, 017–000-121], 1% bovine serum albumin (Sigma-Aldrich,), 1% glycine (Sigma-Aldrich, G5417), 0.2% Triton X-100 for 1 h at room temperature (RT). The Click-iT Plus EdU cell proliferation kit for Imaging, Alexa Fluor 488 dye (Invitrogen, C10637) was then used to tag the integrated EdU according to the manufacture’s protocol. Primary antibodies were diluted in blocking buffer and left to incubate overnight at 4°C. Secondary antibodies, conjugated to Alexa Fluor-fluorochromes, and DAPI for nuclear staining, were diluted 1:300 in blocking buffer and applied for 90 minutes at RT. Glass coverslips were mounted with Fluoromount G to the super plus slides. Imaging was done with the Zeiss LSM880 with Airyscan.

U2OS cells were transfected with siRNAs and incubated in medium containing 20 µM 5-ethynyl-2′-deoxyuridine (EdU, ThermoFisher Scientific, cat# E10187) for 20 minutes, washed and fixed in 2% formaldehyde in PBS for 20 minutes at room temperature. Samples were taken 4 or 6 days after siRNA transfection, depending on the assay.

Next, cells were permeabilized with 0.25% Triton X-100 in PBS for 5 min at room temperature, followed by incubation with click-iT reaction buffer; 2 μM Alexa Fluor 555 or 647 Azide (ThermoFisher Scientific cat# A20012 or A10277), 100 mM TRIS-HCl pH 8.5, 100 mM sodium-L-ascorbate, and 1 mM CuSO4 for 45 min in RT. Cells were then rinsed with PBS, incubated with 0.1 μg/mL4’,6-Diamidino-2-Phenylindole Dihydrochloride (DAPI) and mounted with Aqua Polymount (Polysciences). DNA synthesis was measured by mean EdU intensity / nucleus in replicating cells. Images were acquired using a Zeiss AxioImager M2 widefield fluorescence microscope at 63x magnification (PLAN APO, 1.4 NA, oil-immersion objectives) and were recorded with ZEN 2012 (Blue edition, v1.1.0.0, Zeiss). EdU signal in the nucleus was background corrected. Mean nuclear intensity analysis was done by using Image J.

#### Flow cytometry assessment of EdU incorporation in organoids

For analysis of DNA synthesis via EdU integration assessed by flow cytometry, live organoids were pulsed for 4 hours with 20 uM EdU and then washed once in room temperature DPBS. Organoids then were dissociated with Gentle Cell Dissociation Reagent (Stemcell Technologies, 100-0485) for 10 minutes at 37°C with 5% CO2 and then incubated as a single cell suspension with LIVE/DEAD Fixable Near IR (780) Viability Kit, for 633 nm excitation (Life Technologies, L34992) for 30 minutes on ice to stain dead cells. Cells were washed one time in cold 0.5% BSA PBS and fixed in cold 4% PFA on ice for 15 minutes and then permeabilized with Click-iT saponin-based permeabilization and wash reagent (Invitrogen, C10633) for 30 minutes. The Click-iT Plus EdU Alexa Fluor 488 Flow Cytometry Assay Kit (Invitrogen, C10633) was then used to tag the integrated EdU according to the manufacture’s protocol. Cells were washed three times in cold 0.5% BSA PBS and stained with Alexa Fluor 647 anti-SOX2 (Biolegend, 656108) for 30 minutes on ice. Cells were washed three times in cold 0.5% BSA PBS, filtered, and analyzed with flow cytometer (Beckman Coulter Gallios - 10 color) (Supplementary Fig. 1C). Samples were run in triplicate with 2 organoids per FACS tube across 2 independent differentiations. Only the single, live, SOX2+ cells were analyzed for EdU integration.

#### Immunofluorescence assessment of EdU incorporation in organoids

Organoids were pulsed for 4 hours with 20 μM EdU and then washed three times in room temperature DPBS and then fixed with 4% paraformaldehyde, 4% sucrose (v:v) in PBS (Sigma-Aldrich, P5493) for 30 min. Organoids were washed twice in PBS and transferred to a 30% sucrose solution overnight at 4°C. Organoids were then embedded in Tissue-Tek OCT compound (Sakura Finetech Europe, 4583) and frozen in liquid nitrogen. OCT blocks containing organoids were stored at -80°C prior to sectioning. Organoids were sectioned using a cryostat (Leica) at 20 microns and mounted to charged superfrost plus microscopy slides (Menzel-Glaser, VWR, 900226). After cryosectioning and mounting, organoid sections were outlined on the slides with a hydrophic barrier PAP pen (Sigma, Z377821-1EA). Aspecific antibody binding was blocked by incubation in blocking buffer (PBS, 5% normal goat serum [Invitrogen, 10000 C], 5% normal horse serum [Gibco, 26050088], 5% normal donkey serum [Jackson Immuno Research, 017–000-121], 1% bovine serum albumin (Sigma-Aldrich, A7906), 1% glycine (Sigma-Aldrich, G5417), 0.2% Triton X-100 for 2 h at room temperature (RT). The Click-iT Plus EdU cell proliferation kit for Imaging, Alexa Fluor 488 dye (Invitrogen, C10637) was then used to tag the integrated EdU according to the manufacture’s protocol.

### scEdU-seq on KANSL-depleted cells

U2OS cells were transfected with siLuciferase, siKANSL1 and siKANSL3 and treated with a double pulse of EdU using biological replicate^33^. Samples were made 4 days after treatment with the indicated siRNAs. Azide-PEG3-Biotin was clicked onto EdU labeled DNA as previously described. Subsequently, cells were stained with DAPI to visualize cell cycle progression and labeled with CellTrace for each transfection condition (i.e., mock labeling for siLuciferase, CTY for siKANSL1 and CTFR for siKANSL3) and were sorted (BD Influx) into 384-well plates for scEdU-seq processing. Library preparation included the following steps: proteinase K digestion, genome digestion with NlaIII, DNA blunt-ending, A-tailing, and adapter ligation with cell barcodes and unique molecular identifiers (UMIs). The pooled single-cell libraries were then bound on Dynabeads MyOne Streptavidin C1 (ThermoFisher Scientific) to capture DNA replication fragments. These fragments were released by heat denaturation and subsequently filled in using the Klenow enzyme. The libraries were amplified through in vitro transcription (IVT), reverse transcription (RT), and PCR, followed by sequencing on an Illumina NextSeq 1000 platform (P3, 2×100 bp). Code for analysis and plotting is available on GitHub: https://github.com/jervdberg/KANSL.

### Sample preparation and mass spectrometry data acquisition

Samples preparation for mass spectrometry was described previously^77^. For mass spectrometry, U2OS Flp-In/T-Rex cells expressing GFP-NLS and GFP KANSL3WT were cultured. Cell pellets were lysed in EBC-1 buffer (50 mM Tris, pH 7.5, 150 mM NaCl, 0.5% NP-40, 2 mM MgCl2, protease inhibitor cocktail tablets) with 500 units benzonase. Samples were incubated for 1 hour at 4°C under constant mixing followed by high-speed centrifugation for 10 minutes at 4°C. Protein concentration was measured by Qubit in the cleared lysates, equalized and transferred to tubes containing GFP-Trap beads (Chromotek). After 90 minutes of incubation at 4°C under rotating condition, the beads were washed 4 times with EBC-2 buffer (50 mM Tris pH 7.5, 150 mM NaCl, 1 mM EDTA, and protease inhibitor cocktail tablets) and 3 times with 50 mM ammonium bicarbonate followed by overnight digestion using 2.5 μg trypsin at 37°C under constant shaking. Digestion was terminated with 1% trifluoroacetic acid and centrifuged for 5 minutes at high speed to precipitate insoluble fractions.

Analysis of digested peptides was performed by LC-MS/MS with an Easy-nLC 1000 HPLC system (ThermoFisher) coupled to a Q Exactive Orbitrap mass spectrometer (ThermoFisher). Sample separation was performed on an *in-house* packed 75 µm x 15 cm long silica emitter with a 1.9 µm C18-AQ matrix (Dr. Maisch, Germany) using a 70 minute gradient from 2 – 30 % acetonitrile 0.1% formic acid with a flow rate of 200 nl/minute. A MS survey scan was obtained using a top 10 method for the m/z range of 300-1600, with a maximum injection time of 250 ms, a resolution of 70,000 and an AGC target of 3×10^6^. An isolation mass window of 2.2 m/z was used for the precursor ion selection and the MS/MS Spectra Acquisition was obtained using Higher Energy Collisional Dissociation (HCD) with a normalized collision energy of 25, maximum injection time was 60 ms, with a resolution of 17,500 and an AGC target of 1×10^5^. 20 seconds duration was used for the dynamic exclusion.

### Mass spectrometry data analysis

Mass Spectrometry raw data was analysed using MaxQuant (v1.5.3.30) using standard settings^80^ with the following modifications:. Label Free Quantification was activated. The search was performed against an in silico Trypsin/P-digested *Homo sapiens* proteome (Uniprot May 27th 2019) with a maximum number of mis-cleavages of 4. Match-between-runs with default settings was activated. Maxquant output was further processed in the Perseus computational (v1.6.7.0)^81^. LFQ intensity values were log2-transformed and contaminants,

reverse peptides and proteins identified only by site were removed. Proteins not identified in at least 4 replicates in at least one group were also removed. Missing values were imputed from a normal distribution with a downshift of 1.8 and with of 0.3. Two-tailed t-test were performed to perform statistical comparisons between groups. Perseus statistical output was exported into MS Excel 365 for comprehensive data browsing and visualization. Both raw and processed mass spectrometry data were deposited in the Protein Identification Database (PRIDE)^82^.

### DNA fiber analysis

DNA fibers were prepared as described previously^77, 83^, with minor modifications. In short, both U2OS and iPSCs were labeled with 25 µM 5-chloro-2-deoxyuridine (CldU; Sigma-Aldrich, cat# C6891) for 20 minutes, followed by labeling with 250 µM 5-iodide-2-deoxyuridine (IdU; Sigma-Aldrich, cat# I7125). Subsequently, labeled cells were washed three times in ice cold PBS, harvested and resuspended in cold PBS. Cell concentrations differed between U2OS, iPSCs and U2OS GFP or GFP-RNaseH1 cells, with 2.5×10^5^, 4.0 x10^5^ and 1.0 x10^6^ cell/ml respectively. 2 µl of cell suspension were spotted on a positively charged slide (VWR, cat# 631-0108) and air dried for 5 min. Cells were mixed with 7 µl of lysis buffer (200 mM Tris-HCl pH7.4; 50 mM EDTA; 0.5% SDS), incubated for 2 min, and then tilted to ±45° angle to allow the drop to run by gravity. The slides were air-dried, fixed in methanol/acetic acid (3:1), dried again and stored at RT.

Slides were washed twice in MQ, denatured with 2.5M HCl for 80 min at RT and washed 3x in 1x PBS. Fibers were washed 1x in blocking buffer (1% BSA, 0.1% Tween 20 in PBS) and then incubated in blocking buffer for 20 min. Slides were incubated with 1:500 rat anti-BrdU (clone BU 1/75, Supplementary Table 2) to detect CldU and with 1:750 mouse anti-BrdU (clone B44, Supplementary Table 2) to detect IdU in blocking buffer for 1.5h in the dark. Subsequently, fibers were washed 3 times in 1x PBS, fixed in 2% formaldehyde for 10 min at RT and washed 3 times in 1x PBS. The microscope slides were then incubated with Alexa Fluor 488 donkey anti-rat antibody (1:1000) and Alexa Fluor 555 goat anti-mouse antibody (1:1000) in blocking buffer for 1 h at room temperature in the dark (Supplementary Table 2). Next, slides were washed 4x in 1x PBS and mounted with Aqua Polymount (Polysciences). Samples dried at RT in the dark o/n and subsequently stored at 4°C in the dark until imaging. DNA fibers were visualized and imaged using a Zeiss AxioImager M2 widefield fluorescence microscope at 63x magnification (PLAN APO, 1.4 NA, oil-immersion objectives) and were recorded with ZEN 2012 (Blue edition, v1.1.0.0, Zeiss). Fibers were analyzed using Image J.

Replication fork speed was calculated as kb/min, based on the assumption that 1 µm DNA fiber equals 2.59 kb^84^. Fork symmetry was obtained from the fork speed assay, by analyzing the symmetry between the IdU tracts of bidirectional forks. For fork degradation, cells were incubated with CldU for 20 min, followed by 20 min IdU chase and lastly 4 mM hydroxyurea treatment for 5 h, before trypsinization and slide preparation. IdU track length was divided over CldU tract length, to determine fork degradation. For fork recovery, cells were labelled with CldU for 20 min, followed by 2 h 4 mM hydroxyurea treatment and 20 min IdU labelling. The total amount of fibers was counted, and the percentage of restarted, stalled and new origin fibers determined.

### Immunofluorescence microscopy

iPSCs were plated as single cells at a density of 150 cells/mm^2^ in u-plate 96 well square glass bottom plates (ibidi GmbH, 89627) and fixed after 48 hours for microscopy analysis. Ngn2+ iPSCs differentiated into iNeurons were plated at a density of 100 cells/mm^2^ on nitric acid treated glass cover slips. iPSCs and iNeurons were fixed with 4% paraformaldehyde, 4% sucrose (v:v) in PBS (Sigma-Aldrich, P5493) for 10 min on ice. Aspecific binding was blocked by incubation in brain blocking buffer (PBS, 5% normal goat serum [Invitrogen, 10000 C], 5% normal horse serum [Gibco, 26050088], 5% normal donkey serum [Jackson Immuno Research, 017–000-121], 1% bovine serum albumin (Sigma-Aldrich, A7906), 1% glycine (Sigma-Aldrich, G5417), 0.2% Triton X-100 for 1 h at room temperature (RT). Primary antibodies were diluted in blocking buffer and left to incubate overnight at 4°C. Secondary antibodies, conjugated to Alexa Fluor-fluorochromes (Invitrogen), were diluted 1:300 in blocking buffer for 1 hour at RT, DAPI was added (1:40) for to visualize the nuclei. All antibodies are listed in Supplementary Table 2. For iPSCs, 96 well plate wells were coated with Fluoromount G (Southern Biotech, 0100-01) and for iNeurons, coverslips were mounted with Fluoromount G to glass slides (VWR, 631-1553). The number of functional synapses were quantified per individual cell via manual counting of the co-localized of PSD95 and synapsin puncta and then divided by the dendritic length of the neuron, only primary dendrites were assessed.

Cells were plated on coverslips and treated with siRNAs. Samples were taken 48-72 hours after siRNA treatment, unless otherwise indicated. Next, cells were washes twice in ice cold 1x PBS pre-extracted with 0.5% Triton-X100 in PBS on ice. After washing, cells were fixed with 2% formaldehyde in PBS for 20 min at RT, followed by washing 3x with PBS. Samples were stored o/n at 4°C. For staining, cells were permeabilized with 0.25% Triton for 5 min. at RT. Samples were washed thrice in PBS, blocked in blocking buffer (0.5% BSA, 20mM glycine in 1x PBS) for 20 min. and incubated with primary antibody in blocking buffer for minimal 1.5 h in the dark at RT. Samples were washed twice with blocking buffer and incubated with secondary antibodies (1:1500) and 0.1 μg/mL 4′, 6-diamidino-2-phenylindole dihydrochloride (DAPI) in blocking buffer for 1h in the dark at RT. All antibodies are listed in Supplementary Table 2. Cells were washed with PBS and mounted with Aqua Polymount (Polysciences). Samples were imaged using a Zeiss AxioImager M2 widefield fluorescence microscope at 63x magnification (PLAN APO, 1.4 NA, oil-immersion objectives) and were recorded with ZEN 2012 (Blue edition, v1.1.0.0, Zeiss). Analysis was done in Image J.

Organoids were fixed with 4% paraformaldehyde, 4% sucrose (v:v) in PBS (Sigma-Aldrich, P5493) for 30 min at room temperature. Organoids were washed twice in PBS and transferred to a 30% sucrose solution overnight at 4°C and then embedded in Tissue-Tek OCT compound (Sakura Finetech Europe, 4583) and frozen in liquid nitrogen. OCT blocks containing organoids were stored at -80°C prior to sectioning. 20 μm sections (cryostat, Leica) were mounted to charged superfrost plus microscopy slides (Menzel-Glaser, VWR, 900226). After cryosectioning and mounting, organoid sections were outlined on the slides with a hydrophic barrier PAP pen (Sigma, Z377821-1EA). Aspecific antibody binding was blocked by incubation in brain blocking buffer for 2 h at room temperature (RT). Primary antibodies were diluted in brain blocking buffer and left to incubate overnight at 4°C. Secondary antibodies, conjugated to Alexa Fluor-fluorochromes, and DAPI for nuclear staining, were diluted 1:300 in blocking buffer and applied for 90 minutes at RT. Glass coverslips were mounted with Fluoromount G to the super plus slides. Imaging was done with the Zeiss LSM880 with Airyscan. Signals were background corrected and quantified using FIJI software.

### High throughput imaging of EdU and γH2AX

Cells were grown either on 12 mm sterile glass coverslips or in 96 well plates (ibidi) and transfected with siRNA using Lipofectamine RNAiMAX (ThermoFisher Scientific). Six days after siRNA transfection, cells were labelled with 10 μM of EdU (5-ethynyl-2′-deoxyuridine, ThermoFisher Scientific) for 20 minutes and then fixed with 3% formaldehyde in PBS for 15 minutes at room temperature. The Click-iT EdU Alexa Fluor Imaging Kit (ThermoFisher Scientific) was used for EdU detection. Immunofluorescence was performed as previously described^85^. Briefly, cells were permeabilized for 5 minutes with 0.2 % Triton X-100 (Sigma-Aldrich) in PBS, followed by 3 washes with PBS. Primary and secondary antibodies were diluted in filtered DMEM containing 10 % FBS and 0.02 % sodium azide, and primary antibody incubations were performed for 2 hours at room temperature, whereas secondary antibody incubations were performed for 1 hour at room temperature, with 3 PBS washes performed after each antibody incubation. Cells were then incubated for 10 minutes with PBS containing 4′,6-Diamidino-2-Phenylindole Dihydrochloride (DAPI, 0.5 μg/ml) at room temperature to stain DANN, followed by 3 washes with PBS. The following primary antibody was used in these experiments: H2AX Phospho S139 (mouse, Biolegend 613402, 1:1000). The following secondary antibody was used in these experiments: Alexa Fluor 488 Goat Anti-Mouse (Life Technologies, A11029, 1:500).

### Quantitative image-based cytometry

High-content microscopy for quantitative image-based cytometry (QIBC) was performed as described previously^85^) on a widefield Olympus ScanR High-Content Screening Station (Olympus ScanR Image Acquisition Software version 3.3.0) equipped with an inverted motorized Olympus IX83 microscope, a motorized stage, IR-laser hardware autofocus, a fast emission filter wheel with one set of bandpass filters for multi-wavelength acquisition (DAPI (ex BP 395/25, em BP 435/26), FITC (ex BP 470/24, em BP 511/23), TRITC (ex BP 550/15, em BP 595/40), Cy5 (ex BP 640/30, em BP 705/72)), and a Hamamatsu ORCA-FLASH 4.0 V2 sCMOS camera (2048 × 2048 pixel, pixel size 6.5 × 6.5 μm) with a 20x UPLSAPO (NA 0.75) air objective. All images were acquired under non-saturating conditions, and identical settings were applied to all samples within one experiment. Images were analyzed with the Olympus ScanR Image Analysis Software (version 3.3.0). A dynamic background correction was applied, and nuclei segmentation was performed using an integrated intensity-based object detection module based on the DAPI signal. Downstream analyses were focused on properly detected interphase nuclei containing a 2N-4N DNA content as measured by total and mean DAPI intensities. Fluorescence intensities are depicted as arbitrary units.

### R-loop detection via immunofluorescence microscopy

R-loops were detected via two different methods; via S9.6 antibody imaging (with or without GFP-RNaseH1 WT protein incubation) and via GFP-RNaseH1 D210N imaging. For both assays cells are seeded in a way to assure equal cell density at the moment of fixation.

For imaging via the S9.6 antibody, siRNA treated cells were fixed via 2% formaldehyde or via methanol. In the latter case, cells were washed twice in 1x PBS and subsequently are fixed in 100% ice cold methanol for 10 min at -20°C. After this, the cells are washed 5x in 1x PBS and stored o/n at 4°C. After fixation, S9.6 staining is performed via immunofluorescence protocol described above, using the S9.6 antibody and anti-nucleolin antibody (Supplementary Table 2).

When samples were treated with GFP-RNaseH1 WT protein, samples were pre-extracted with 0.5% ice cold Triton-X100 in PBS, fixed in 2% formaldehyde and stored o/n at 4°C. Cells were post-extracted with 0.25% Triton-X100 in PBS and washed thrice. Samples were washed in 1x RNaseH1 buffer, and GFP-RNaseH1 WT protein (1:1800, 0.188 mg/ml) was incubated in 1x RNaseH1 buffer for 4h at 37°C. After this, the samples were washed thrice and the staining continued with the S9.6 staining described above, with the only deviation being that primary antibody incubation occurred over night. Samples were imaged using a Zeiss AxioImager M2 widefield fluorescence microscope at 63x magnification (PLAN APO, 1.4 NA, oil-immersion objectives) and were recorded with ZEN 2012 (Blue edition, v1.1.0.0, Zeiss). Analysis was done in Image J, where the S9.6 signal of the nucleoli is excluded from the rest of the nuclear signal.

Staining with GFP-RNaseH1 D210N was done as described previously^44^, with minor modifications. Samples were pre-extracted with 0.5% ice cold Triton-X100 in PBS, fixed in 2% formaldehyde and stored o/n at 4°C. Cells were post-extracted with 0.25% Triton-X100 in PBS and washed thrice. Samples were washed in 1x RNaseH1 buffer, and GFP-RNaseH1 D210N protein (1:1800, 0.188 mg/ml) was incubated in 1x RNaseH1 buffer for 4h at 37°C. After this cells were incubated with DAPI (0.1 μg/mL) in PBS for 30 min. in PBS in the dark, and after wa shing mounted with Aqua Polymount (Polysciences). Samples were imaged using a Zeiss AxioImager M2 widefield fluorescence microscope at 63x magnification (PLAN APO, 1.4 NA, oil-immersion objectives) and were recorded with ZEN 2012 (Blue edition, v1.1.0.0, Zeiss). Analysis was done in Image J, and the mean GFP intensity per nucleus was assessed.

### Lentiviral GFP and GFP-RNaseH1 treatment

HEK293T cells were transfected with GFP or GFP-RNaseH1, using PEI (1:6 ratio to DNA). After transfection the medium of the cells was replaced with DMEM + 10mM Hepes. The virus containing medium was harvested 24-48h later, centrifuged and the supernatant was filtered before snap freezing the virus and storing it at -80°C. Virus transduction was performed on U2OS cells, with a MOI > 1. Polybrene (4 µg/ml) was added to improve viral transduction. The GFP and GFP-RNaseH1 constructs were a gift from the Luijsterburg lab (LUMC, the Netherlands). Rescue experiments with GFP and GFP-RNaseH1 in U2OS cells were performed as followed: Cells were reverse transfected with siRNAs and incubated for ±6h at 37°C. During seeding more cells were seeded for NSL-depleted samples, to correct for NSL-dependent reduced proliferation. After incubation, cells were transduced with either GFP or GFP-RNaseH1. Samples were harvested 4 days after transfection and transduction, and EdU incorporation or fork symmetry was assessed as described above.

### High-content Proximity Ligation Assay (PLA)

PLA experiments were performed as previously described with minor modifications^86^. Briefly, siRNA transfected cells were grown on coverslips and pulse-labeled with 10 µM EdU for 30 minutes. Slides were washed with ice-cold 4X Saline-sodium citrate (SSC) buffer and fixed in -20°C methanol for 10 minutes. After washing with ice-cold SSC, slides were incubated with acetone for 1 minute on ice. Slides were then washed three times with ice-cold SSC buffer, and incubated for 10 minutes in ice-cold SSC buffer. Slides were blocked with blocking buffer (3% BSA in SSC with 0.1%Tween) for 1 hour at room temperature and washed once with PBS. Freshly prepared Click-iT reaction mix (2mM copper sulfate, 10µM biotin-azide and 100mM sodium ascorbate in PBS) was added to all samples at room temperature for 1 hour. Slides were incubated with primary antibodies at 4°C overnight (S9.6 antibody 1:50^87^, Anti-biotin antibody 1:500). Following the manufacturer’s protocol, slides were washed three times using buffer A (0.01M Tris, 0.15M NaCl and 0.05% Tween-20, pH 815 7,4) for 5 minutes each. Subsequently, slides were incubated with Duolink in Situ PLA probes anti-mouse plus and anti-rabbit minus for 1 hour at 37°C in a humid chamber. After washes with buffer A, slides were incubated with Duolink ligation mix for 30 minutes at 37°C in a humid chamber. Slides were washed twice in buffer A for 2 minutes each and then incubated with Duolink amplification mix for exactly 120 minutes at 37°C. Slides were washed three times with buffer B (0.2M Tris and 0.1M NaCl in PBS) for 10 minutes each and one time with 0.01x buffer B. DNA was stained using DAPI for 15 minutes and slides were mounted with ProLongTM Gold antifade mount (Invitrogen). Slides were kept in dark at 4°C until imaged. Images were captured using Metafer5 and quantified using MetaSystem.

### Clonogenic survival assays

Cell survival assays were performed as described previously^77^, with some minor variations. HCT116 cells were transfected with siRNAs, trypsinized, seeded at low density and exposed /to HU for 24 hours. For HCT116 Flp-In/T-Rex, cDNAs were expressed by adding doxycycline for 24 hours after siRNA transfection. After 8-12 days, the cells were washed with 0.9% NaCl and stained with methylene blue (2.5 g/L in 5% ethanol, Sigma-Aldrich). Colonies of more than 20 cells were scored. Analysis was done via Image J.

### DRIP-seq

DRIP-seq was performed as previously described^88^ with minor modifications. Cells were transfected with siLuc or siKANSL3 as described previously. Cells were harvested 4 days after siRNA transfection. Samples were sequenced using an Illumina NextSeq500 or HiSeq X, using paired-end sequencing with 42 or 151 bp from each end.

### DRIP-seq data analyses

For all sequencing data, a sequencing quality profile was generated using FastQC (Version 0.11.9). Sequences were trimmed using TrimGalore (Version 0.6.5) and Cutadapt (version 2.9). Reads were aligned to the Human Genome 38 using bwa-mem tools (BWA (Version 0.7.17) and reference genome GCA_000001405.15_GRCh38^89^. Only uniquely mapping and high-quality reads (> q30) were included in the analyses. Bam files were converted into stranded TagDirectories and genome tracks using Hypergeometric Optimization of Motif EnRichment tools (HOMER Version 4.8.2; with forced fragment lengths of 150bp)^90^. Example genome tracks were generated in Integrative Genomics Viewer (IGV; version 2.4.3). A list of 49,948 gene-coordinates was obtained from the University of California Santa Cruz (UCSC) genome database selecting the “knownCanonical” table containing the canonical transcription start sites per gene^91^. Read-density profiles around TSS coordinates were defined using the AnnotatePeaks.pl tool of HOMER using 10bp bins and the default normalization to 10 million reads. Individual datasets were subsequently processed into heatmaps or binding profiles using R (Version 4.0.5) and Rstudio (Version 1.1.423)^92^. To define TSSs with increased/ decreased DRIP signal in siKANSL3-treated cells, we counted the number of reads around each TSS (+/- 1kb) and performed t-test comparisons between the three siLuc and three siKANSL3 repeats, using p=0.05 as significance cutoff. To prevent contamination of binding profiles by small genes, we included only genes of at least 5kb in size.. PeakCalling over the input was performed for all individual siLuc and siKANSL3-treated DRIP-seq samples, using the findPeaks tool from HOMER. Only peaks identified in all three siLuc or all three siKANSL3 repeats were extracted using the HOMER tool mergePeaks with -d given. The remaining peaks in siLuc and siKANSL3 were again merged using HOMER tool mergePeaks with -d given and all peaks identified in either siLuc, siKANSL3 or both, were included in the downstream analyses.

#### ChIP-seq datasets downloaded from literature

H4K5ac and H4K8ac ChIP-seq data were downloaded from GSE158736^19^. RNAPII ChIP-seq data were obtained from GSE140930 (GSM4190607)^50^. Datafiles were trimmed and mapped to hg38 as described for DRIP-seq. For the RNAPII ChIP-seq datasets duplicate reads were removed using Samtools (Version 1.6) with fixmate -m and markdup -r settings. Bam files were converted into stranded TagDirectories and UCSC genome tracks using HOMER tools (Version 4.8.2) and further analyzed as described for the DRIP-seq data.

### TTchem-seq

Nascent RNA-seq (TTchem-seq) was performed largely as previously described^51^. Cells were prepared in a similar fashion as for the DRIP-seq. On day 4 after siRNA transfection,cells were labelled with 1 mM 4SU (Glentham Life Sciences) for 30 minutes. 4TU-labeled (Sigma) yeast spike-in RNA was generated as described in^51^. Immediately after 4SU labelling, cells were washed with PBS and lysed in TRIzol. 0.8 µg spike-in RNA was added per 10 million cells, resulting in addition of ∼0.25% spike RNA per condition. RNA was isolated by TRIzol– chloroform isolation and ethanol precipitation. Then, 400 μg RNA (in a total volume of 116 μl) was fragmented by adding 4 μl 5 M NaOH and incubating on ice for 30 min, then stopped by addition of 80 μl of 1 M Tris pH 7 and cleaned up twice using the Micro Bio-Spin P-30 Gel Columns (Bio-Rad, 7326223) according to the manufacturer’s instructions. Biotinylation of 4SU residues was performed in a total volume of 250 μl, containing 10 mM Tris–HCl pH 7.4, 1 mM EDTA and 5 µg of MTSEA biotin-XX linker (Biotium, BT90066) for 30 min at RT in the dark, followed by phenol–chloroform extraction and elution in 50ul water. 300 µg RNA (37.5 µl) was denatured by 10 min incubation at 65 °C and added to 100 μl μMACS Streptavidin MicroBeads (Milentyl, 130-074-101). RNA was incubated with beads for 15 min at RT and beads were applied to a μColumn in the magnetic field of a μMACS magnetic separator. The beads were washed three times with 55 °C pulldown wash buffer (100 mM Tris–HCl pH 7.4, 10 mM EDTA, 1 M NaCl and 0.1% Tween-20). Biotinylated RNA was eluted twice by addition of 100 mM DTT and cleaned up using the RNeasy MinElute kit (Qiagen, 74204) using 1,050 μl ethanol (≥99%) per 200 μl reaction after addition of 700 μl of RLT buffer to precipitate RNA of less than 200 nucleotides. 500 to 800 ng of the purified 4SU-labelled RNA was then used as input for in-house stranded RNA library prep, similar to the NEBNext® Ultra™ II Directional RNA Library Prep Kit for Illumina (NEB #E7760) without rRNA depletion. The library was amplified with ten PCR cycles and quality-control checked on the TapeStation (Agilent) using the High Sensitivity DNA Kit before pooling and paired-end sequencing on the NovaseqX system.

### TTchem-seq data analyses

A sequencing quality profile was generated using FastQC (version 0.11.9). If needed, sequences were trimmed using TrimGalore (version 0.6.5). A combined genome was generated, containing genome GCA_000001405.15_GRCh38_no_alt_analysis_set and Saccharomyces_cerevisiae.R64-1-1.dna.toplevel (and sjdbGTFfile Homo_sapiens.GRCh38.94.correctedContigs.gtf and Saccharomyces_cerevisiae.R64-1-1.113.gtf). Reads were aligned to the combined genome using STAR (version 2.7.7a/gcc-8.3.1). Duplicate reads were removed using Samtools (Version 1.11) with fixmate -m and markdup -r settings. Reads mapping to the S cerevisiae genome were used for spike-in correction of downstream analyses. Bam files were converted into stranded TagDirectories and spike-normalized genome tracks using HOMER tools (Version 4.8.2) with forced fragment lengths of 150 bp^93^. For a first genome-wide comparison of transcription levels between WT and KANSL-depleted cells, a list of 9,944 gene-coordinates was obtained from the University of California Santa Cruz (UCSC) genome database, selecting the “knownCanonical” table containing the canonical transcription start sites per gene, and further selecting for only non-overlapping genes of over 3kb with at least 2kb between genes^91^. For this list of genes, TTchem-seq levels were quantified in 200bp bins from -5kb to +100kb around the TSS, using the AnnotatePeaks.pl tool of HOMER, and subsequently normalized to spike-in reads. Independently, more specific read-densities around DRIP peaks or TSSs with significantly increased or decreased DRIP signal in siKANSL3 versus siLuc - identified as described in the section on DRIP-seq analyses - were defined per 10bp bins and also normalized to spike-in reads. Read-densities around TSSs or DRIP peaks - identified as described in the section on DRIP-seq analyses - were defined per 10bp bins using the AnnotatePeaks.pl tool of HOMER and subsequently normalized to spike-in reads. Individual datasets were subsequently processed into heatmaps or binding profiles using R (Version 4.0.5) and Rstudio (Version 1.1.423)^92^.

### Statistical analysis

Statistical of the data was performed using GraphPad Prism 10 (GraphPad Software, Inc., CA, USA).

## Acknowledgements

The authors thanks Magda Rother, Colette Moses, Frank Jacobs, and Sofia Puvogel for insightful discussions, and Anton de Groot, Chantal Schoenmaker, Astrid Oudakker, Denise Duineveld, and Ka Man Wu for their experimental support. This work was financially supported by grants from the European Research Council (ERC-AdG, 101053581-scTranslatomics to A.v.O; ERC-CoG, 101043815-STOP-FIX-GO to M.S.L), the Novo Nordisk Fonden Synergy Programme (Lost Memories, 0091873 to J.v.d.B. and A.v.O.), the Swiss National Science Foundation (310030-197003 to M.A.). the Dutch Research Council (NWO ENW-M2, OCENW.M20.216 to N.N.K. and B.L.L., NWO-VICI, 182.052 to H.v.A.), the Simons Foundation Autism Research Initiative (SFARI, 890042 to N.N.K.), and the Koolen-de Vries Foundation (to B.L.L.).

## Author contributions

J.L. performed immunofluorescence, western blot, RT-qPCR and DNA fiber analysis, assisted in scEdU-seq, DRIP-seq and TTchem experiments and wrote the paper. D.v.d.H. performed TTchem-seq and analyzed DRIP-seq data. J.v.d.B. performed scEdu-seq, analyzed the data and wrote the paper. J.K.S. generated stable cell lines and performed immunofluorescence analysis and clonogenic survivals. R.v.H. performed immunofluorescence, western blot and RT-qPCR analysis and clonogenic assays. W.W. performed DNA fiber and cell proliferation assays. D.E.C.B. performed DRIP-seq. C.S.Y.L. performed PLA assays. A.P. performed immunofluorescence and high-throughput imaging. R.G-P. performed mass spectrometry analysis. M.F. performed immunofluorescence analysis. N.K. performed iNeuron experiments.

O.G.d.J. performed plasmid design and cloning. A.C.O.V. supervised mass spectrometry analysis. M.A. supervised high-throughput imaging and analysis. N.T. supervised PLA assays.

A.v.O. supervised scEdu-seq experiments. M.S.L. supervised DRIP-seq and TTchem-seq experiments, analyzed the data and wrote the paper. N.N.K. supervised the project. B.L.L. performed immunofluorescence, flow cytometry and western blot analysis, proliferation assays, organoid and iNeuron experiments, and wrote the paper. H.v.A. supervised the project and wrote the paper. All authors commented on and approved the manuscript

## Additional information

**Extended Data** is available for this paper in the Extended Data file.

**Supplementary information** is available for this paper in the Supplementary Information file.

## Correspondence and Materials availability

Correspondence and requests for materials should be addressed to Haico van Attikum and Brooke L. Latour.

## References

1. Qing, X., Zhang, G. & Wang, Z.Q. DNA damage response in neurodevelopment and neuromaintenance. Febs j 290, 3300–3310 (2023).

2. McConnell, M.J. et al. Intersection of diverse neuronal genomes and neuropsychiatric disease: The Brain Somatic Mosaicism Network. Science 356 (2017).

3. Dehay, C. & Kennedy, H. Cell-cycle control and cortical development. Nat Rev Neurosci 8, 438–450 (2007).

4. Ernst, C. Proliferation and Differentiation Deficits are a Major Convergence Point for Neurodevelopmental Disorders. Trends Neurosci 39, 290–299 (2016).

5. Zeman, M.K. & Cimprich, K.A. Causes and consequences of replication stress. Nat Cell Biol 16, 2–9 (2014).

6. Bellelli, R. & Boulton, S.J. Spotlight on the Replisome: Aetiology of DNA Replication-Associated Genetic Diseases, in Trends Genet, Vol. 37 317–336 (© 2020 The Author(s). Published by Elsevier Ltd., England; 2021).

7. Chakraborty, A. et al. Replication Stress Induces Global Chromosome Breakage in the Fragile X Genome. Cell Rep 32, 108179 (2020).

8. Van Esch, H. et al. Defective DNA Polymerase α-Primase Leads to X-Linked Intellectual Disability Associated with Severe Growth Retardation, Microcephaly, and Hypogonadism. Am J Hum Genet 104, 957–967 (2019).

9. Alderton, G.K. et al. Seckel syndrome exhibits cellular features demonstrating defects in the ATR-signalling pathway. Hum Mol Genet 13, 3127–3138 (2004).

10. Koolen, D.A. et al. A new chromosome 17q21.31 microdeletion syndrome associated with a common inversion polymorphism, in Nat Genet, Vol. 38 999-1001 (United States; 2006).

11. Schmit, M. & Bielinsky, A.K. Congenital Diseases of DNA Replication: Clinical Phenotypes and Molecular Mechanisms. Int J Mol Sci 22 (2021).

12. Koolen, D.A. et al. The Koolen-de Vries syndrome: a phenotypic comparison of patients with a 17q21.31 microdeletion versus a KANSL1 sequence variant. Eur J Hum Genet 24, 652–659 (2016).

13. Myers, K.A. et al. The epileptology of Koolen-de Vries syndrome: Electro-clinico-radiologic findings in 31 patients. Epilepsia 58, 1085–1094 (2017).

14. Zollino, M. et al. Mutations in KANSL1 cause the 17q21.31 microdeletion syndrome phenotype. Nat Genet 44, 636–638 (2012).

15. Koolen, D.A. et al. Mutations in the chromatin modifier gene KANSL1 cause the 17q21.31 microdeletion syndrome. Nat Genet 44, 639–641 (2012).

16. Dias, J. et al. Structural analysis of the KANSL1/WDR5/KANSL2 complex reveals that WDR5 is required for efficient assembly and chromatin targeting of the NSL complex. Genes Dev 28, 929–942 (2014).

17. Cai, Y. et al. Subunit composition and substrate specificity of a MOF-containing histone acetyltransferase distinct from the male-specific lethal (MSL) complex. J Biol Chem 285, 4268–4272 (2010).

18. Sheikh, B.N., Guhathakurta, S. & Akhtar, A. The non-specific lethal (NSL) complex at the crossroads of transcriptional control and cellular homeostasis. EMBO Rep 20, e47630 (2019).

19. Radzisheuskaya, A. et al. Complex-dependent histone acetyltransferase activity of KAT8 determines its role in transcription and cellular homeostasis. Mol Cell 81, 1749–1765 e1748 (2021).

20. Feller, C. et al. The MOF-containing NSL complex associates globally with housekeeping genes, but activates only a defined subset. Nucleic Acids Res 40, 1509–1522 (2012).

21. Zhao, X. et al. Crosstalk between NSL histone acetyltransferase and MLL/SET complexes: NSL complex functions in promoting histone H3K4 di-methylation activity by MLL/SET complexes. PLoS Genet 9, e1003940 (2013).

22. Ravens, S. et al. Mof-associated complexes have overlapping and unique roles in regulating pluripotency in embryonic stem cells and during differentiation. Elife 3 (2014).

23. Lam, K.C. et al. The NSL complex-mediated nucleosome landscape is required to maintain transcription fidelity and suppression of transcription noise. Genes Dev 33, 452–465 (2019).

24. Gaub, A. et al. Evolutionary conserved NSL complex/BRD4 axis controls transcription activation via histone acetylation. Nat Commun 11, 2243 (2020).

25. Linda, K. et al. Imbalanced autophagy causes synaptic deficits in a human model for neurodevelopmental disorders. Autophagy 18, 423–442 (2022).

26. Karoutas, A. et al. The NSL complex maintains nuclear architecture stability via lamin A/C acetylation. Nat Cell Biol 21, 1248–1260 (2019).

27. Singh, D.K. et al. MOF Suppresses Replication Stress and Contributes to Resolution of Stalled Replication Forks. Mol Cell Biol 38 (2018).

28. Li, L. et al. Lysine acetyltransferase 8 is involved in cerebral development and syndromic intellectual disability. J Clin Invest 130, 1431–1445 (2020).

29. Verboven, A.H.A. et al. Integrative transcriptomics and electrophysiological profiling of hiPSC-derived neurons identifies novel druggable pathways in Koolen-de Vries Syndrome. bioRxiv, 2024.2008.2029.610281 (2024).

30. Meunier, S. et al. An epigenetic regulator emerges as microtubule minus-end binding and stabilizing factor in mitosis. Nat Commun 6, 7889 (2015).

31. Rodriguez-Acebes, S., Mourón, S. & Méndez, J. Uncoupling fork speed and origin activity to identify the primary cause of replicative stress phenotypes. J Biol Chem 293, 12855–12861 (2018).

32. Sedlackova, H. et al. Equilibrium between nascent and parental MCM proteins protects replicating genomes. Nature 587, 297–302 (2020).

33. van den Berg, J., et al. Quantifying DNA replication speeds in single cells by scEdU-seq. Nat Methods 21, 1175–1184 (2024).

34. Daigh, L.H., Liu, C., Chung, M., Cimprich, K.A. & Meyer, T. Stochastic Endogenous Replication Stress Causes ATR-Triggered Fluctuations in CDK2 Activity that Dynamically Adjust Global DNA Synthesis Rates. Cell Syst 7, 17–27.e13 (2018).

35. Bianchi, V., Pontis, E. & Reichard, P. Changes of deoxyribonucleoside triphosphate pools induced by hydroxyurea and their relation to DNA synthesis. J Biol Chem 261, 16037–16042 (1986).

36. Petermann, E., Orta, M.L., Issaeva, N., Schultz, N. & Helleday, T. Hydroxyurea-stalled replication forks become progressively inactivated and require two different RAD51-mediated pathways for restart and repair. Mol Cell 37, 492–502 (2010).

37. Zellweger, R. et al. Rad51-mediated replication fork reversal is a global response to genotoxic treatments in human cells. J Cell Biol 208, 563–579 (2015).

38. Schlacher, K. et al. Double-strand break repair-independent role for BRCA2 in blocking stalled replication fork degradation by MRE11. Cell 145, 529–542 (2011).

39. Vassin, V.M., Anantha, R.W., Sokolova, E., Kanner, S. & Borowiec, J.A. Human RPA phosphorylation by ATR stimulates DNA synthesis and prevents ssDNA accumulation during DNA-replication stress. J Cell Sci 122, 4070–4080 (2009).

40. Gaillard, H. & Aguilera, A. Transcription as a Threat to Genome Integrity. Annu Rev Biochem 85, 291–317 (2016).

41. Gómez-González, B. & Aguilera, A. Transcription-mediated replication hindrance: a major driver of genome instability. Genes Dev 33, 1008–1026 (2019).

42. Crossley, M.P., Bocek, M. & Cimprich, K.A. R-Loops as Cellular Regulators and Genomic Threats. Mol Cell 73, 398–411 (2019).

43. Uruci, S., Lo, C.S.Y., Wheeler, D. & Taneja, N. R-Loops and Its Chro-Mates: The Strange Case of Dr. Jekyll and Mr. Hyde. Int J Mol Sci 22 (2021).

44. Crossley, M.P. et al. Catalytically inactive, purified RNase H1: A specific and sensitive probe for RNA-DNA hybrid imaging. J Cell Biol 220 (2021).

45. Cerritelli, S.M. & Crouch, R.J. Ribonuclease H: the enzymes in eukaryotes. Febs j 276, 1494–1505 (2009).

46. Skourti-Stathaki, K., Proudfoot, N.J. & Gromak, N. Human senataxin resolves RNA/DNA hybrids formed at transcriptional pause sites to promote Xrn2-dependent termination. Mol Cell 42, 794–805 (2011).

47. Hamperl, S., Bocek, M.J., Saldivar, J.C., Swigut, T. & Cimprich, K.A. Transcription-Replication Conflict Orientation Modulates R-Loop Levels and Activates Distinct DNA Damage Responses. Cell 170, 774–786.e719 (2017).

48. Perez-Calero, C. et al. UAP56/DDX39B is a major cotranscriptional RNA-DNA helicase that unwinds harmful R loops genome-wide. Genes Dev 34, 898–912 (2020).

49. Goehring, L., Huang, T.T. & Smith, D.J. Transcription-Replication Conflicts as a Source of Genome Instability. Annu Rev Genet 57, 157–179 (2023).

50. van den Heuvel, D., et al. A CSB-PAF1C axis restores processive transcription elongation after DNA damage repair. Nat Commun 12, 1342 (2021).

51. Gregersen, L.H., Mitter, R. & Svejstrup, J.Q. Using TT(chem)-seq for profiling nascent transcription and measuring transcript elongation. Nature protocols 15, 604–627 (2020).

52. Ginno, P.A., Lim, Y.W., Lott, P.L., Korf, I. & Chedin, F. GC skew at the 5’ and 3’ ends of human genes links R-loop formation to epigenetic regulation and transcription termination. Genome research 23, 1590–1600 (2013).

53. Castillo-Guzman, D., Hartono, S.R., Sanz, L.A. & Chédin, F. SF3B1-targeted Splicing Inhibition Triggers Global Alterations in Transcriptional Dynamics and R-Loop Metabolism. bioRxiv, 2020.2006.2008.130583 (2020).

54. Wan, Y. et al. Splicing function of mitotic regulators links R-loop-mediated DNA damage to tumor cell killing. J Cell Biol 209, 235–246 (2015).

55. Liu, Z.S. et al. R-Loop Accumulation in Spliceosome Mutant Leukemias Confers Sensitivity to PARP1 Inhibition by Triggering Transcription-Replication Conflicts. Cancer Res 84, 577–597 (2024).

56. Matos, D.A. et al. ATR Protects the Genome against R Loops through a MUS81-Triggered Feedback Loop. Mol Cell 77, 514–527 e514 (2020).

57. Benito-Kwiecinski, S. et al. An early cell shape transition drives evolutionary expansion of the human forebrain. Cell 184, 2084–2102.e2019 (2021).

58. Lv, X. et al. TBR2 coordinates neurogenesis expansion and precise microcircuit organization via Protocadherin 19 in the mammalian cortex. Nat Commun 10, 3946 (2019).

59. LaMarca, E.A., et al. R-loop landscapes in the developing human brain are linked to neural differentiation and cell-type specific transcription. bioRxiv (2023).

60. Hamperl, S., Bocek, M.J., Saldivar, J.C., Swigut, T. & Cimprich, K.A. Transcription-Replication Conflict Orientation Modulates R-Loop Levels and Activates Distinct DNA Damage Responses. Cell 170, 774–786 e719 (2017).

61. Arbogast, T. et al. Mouse models of 17q21.31 microdeletion and microduplication syndromes highlight the importance of Kansl1 for cognition. PLoS Genet 13, e1006886 (2017).

62. García-Muse, T. & Aguilera, A. R Loops: From Physiological to Pathological Roles. Cell 179, 604–618 (2019).

63. Stork, C.T. et al. Co-transcriptional R-loops are the main cause of estrogen-induced DNA damage. Elife 5 (2016).

64. Wahba, L., Costantino, L., Tan, F.J., Zimmer, A. & Koshland, D. S1-DRIP-seq identifies high expression and polyA tracts as major contributors to R-loop formation. Genes Dev 30, 1327–1338 (2016).

65. Niehrs, C. & Luke, B. Regulatory R-loops as facilitators of gene expression and genome stability. Nat Rev Mol Cell Biol 21, 167–178 (2020).

66. Ginno, P.A., Lott, P.L., Christensen, H.C., Korf, I. & Chédin, F. R-loop formation is a distinctive characteristic of unmethylated human CpG island promoters. Mol Cell 45, 814–825 (2012).

67. Nadel, J. et al. RNA:DNA hybrids in the human genome have distinctive nucleotide characteristics, chromatin composition, and transcriptional relationships. Epigenetics Chromatin 8, 46 (2015).

68. Grunseich, C. et al. Senataxin Mutation Reveals How R-Loops Promote Transcription by Blocking DNA Methylation at Gene Promoters. Mol Cell 69, 426–437.e427 (2018).

69. Castellano-Pozo, M. et al. R loops are linked to histone H3 S10 phosphorylation and chromatin condensation. Mol Cell 52, 583–590 (2013).

70. Groh, M., Lufino, M.M., Wade-Martins, R. & Gromak, N. R-loops associated with triplet repeat expansions promote gene silencing in Friedreich ataxia and fragile X syndrome. PLoS Genet 10, e1004318 (2014).

71. Thongthip, S., Carlson, A., Crossley, M.P. & Schwer, B. Relationships between genome-wide R-loop distribution and classes of recurrent DNA breaks in neural stem/progenitor cells. Sci Rep 12, 13373 (2022).

72. Corazzi, L. et al. Linear interaction between replication and transcription shapes DNA break dynamics at recurrent DNA break Clusters. Nat Commun 15, 3594 (2024).

73. Tingler, M., Philipp, M. & Burkhalter, M.D. DNA Replication proteins in primary microcephaly syndromes. Biol Cell 114, 143–159 (2022).

74. Charlier, C.F. & Martins, R.A.P. Protective Mechanisms Against DNA Replication Stress in the Nervous System. Genes (Basel) 11 (2020).

75. Smits, D.J. et al. De novo MCM6 variants in neurodevelopmental disorders: a recognizable phenotype related to zinc binding residues. Hum Genet 142, 949–964 (2023).

76. Saxonov, S., Berg, P. & Brutlag, D.L. A genome-wide analysis of CpG dinucleotides in the human genome distinguishes two distinct classes of promoters. Proc Natl Acad Sci U S A 103, 1412–1417 (2006).

77. Singh, J.K. Identification and characterization of novel factors in the DNA damage response. Human Genetics, 245. Faculty of Medicine, Leiden University Medical Center (LUMC), Leiden, The Netherlands. Retrieved from https://hdl.handle.net/1887/3485639 (2022).

78. Frega, M. et al. Rapid Neuronal Differentiation of Induced Pluripotent Stem Cells for Measuring Network Activity on Micro-electrode Arrays. J Vis Exp (2017).

79. van de Kooij, B. et al. The Fanconi anemia core complex promotes CtIP-dependent end resection to drive homologous recombination at DNA double-strand breaks. Nat Commun 15, 7076 (2024).

80. Tyanova, S., Temu, T. & Cox, J. The MaxQuant computational platform for mass spectrometry-based shotgun proteomics. Nat Protoc 11, 2301–2319 (2016).

81. Tyanova, S. et al. The Perseus computational platform for comprehensive analysis of (prote)omics data. Nat Methods 13, 731–740 (2016).

82. Perez-Riverol, Y. et al. The PRIDE database at 20 years: 2025 update. Nucleic Acids Res 53, D543–d553 (2025).

83. Nieminuszczy, J., Schwab, R.A. & Niedzwiedz, W. The DNA fibre technique - tracking helicases at work. Methods 108, 92–98 (2016).

84. Jackson, D.A. & Pombo, A. Replicon clusters are stable units of chromosome structure: evidence that nuclear organization contributes to the efficient activation and propagation of S phase in human cells. J Cell Biol 140, 1285–1295 (1998).

85. Lezaja, A. et al. RPA shields inherited DNA lesions for post-mitotic DNA synthesis. Nat Commun 12, 3827 (2021).

86. Gaggioli, V. et al. Dynamic de novo heterochromatin assembly and disassembly at replication forks ensures fork stability. Nat Cell Biol 25, 1017–1032 (2023).

87. De Magis, A. et al. DNA damage and genome instability by G-quadruplex ligands are mediated by R loops in human cancer cells. Proc Natl Acad Sci U S A 116, 816–825 (2019).

88. Sanz, L.A. & Chedin, F. High-resolution, strand-specific R-loop mapping via S9.6-based DNA-RNA immunoprecipitation and high-throughput sequencing. Nat Protoc 14, 1734–1755 (2019).

89. Li, H. Aligning sequence reads, clone sequences and assembly contigs with BWA-MEM. arXiv:1303.3997v2 [q-bio.GN] (2013).

90. Benaglia T, C.D., Hunter DR, Young D mixtools: An R Package for Analyzing Finite Mixture Models. Journal of Statistical Software 32, 1–29 (2009).

91. Karolchik, D. et al. The UCSC Table Browser data retrieval tool. Nucleic Acids Res 32, D493–496 (2004).

92. Team, R.C. R: A language and environment for statistical computing. (2019).

93. Heinz, S. et al. Simple combinations of lineage-determining transcription factors prime cis-regulatory elements required for macrophage and B cell identities. Mol Cell 38, 576–589 (2010).

