## Supplementary material for "The NSL complex promotes neural development by preventing R-loop induced replication stress": Lingeman_et_al_Supplementary_data

Lingeman et al_Extended Data Figure 1

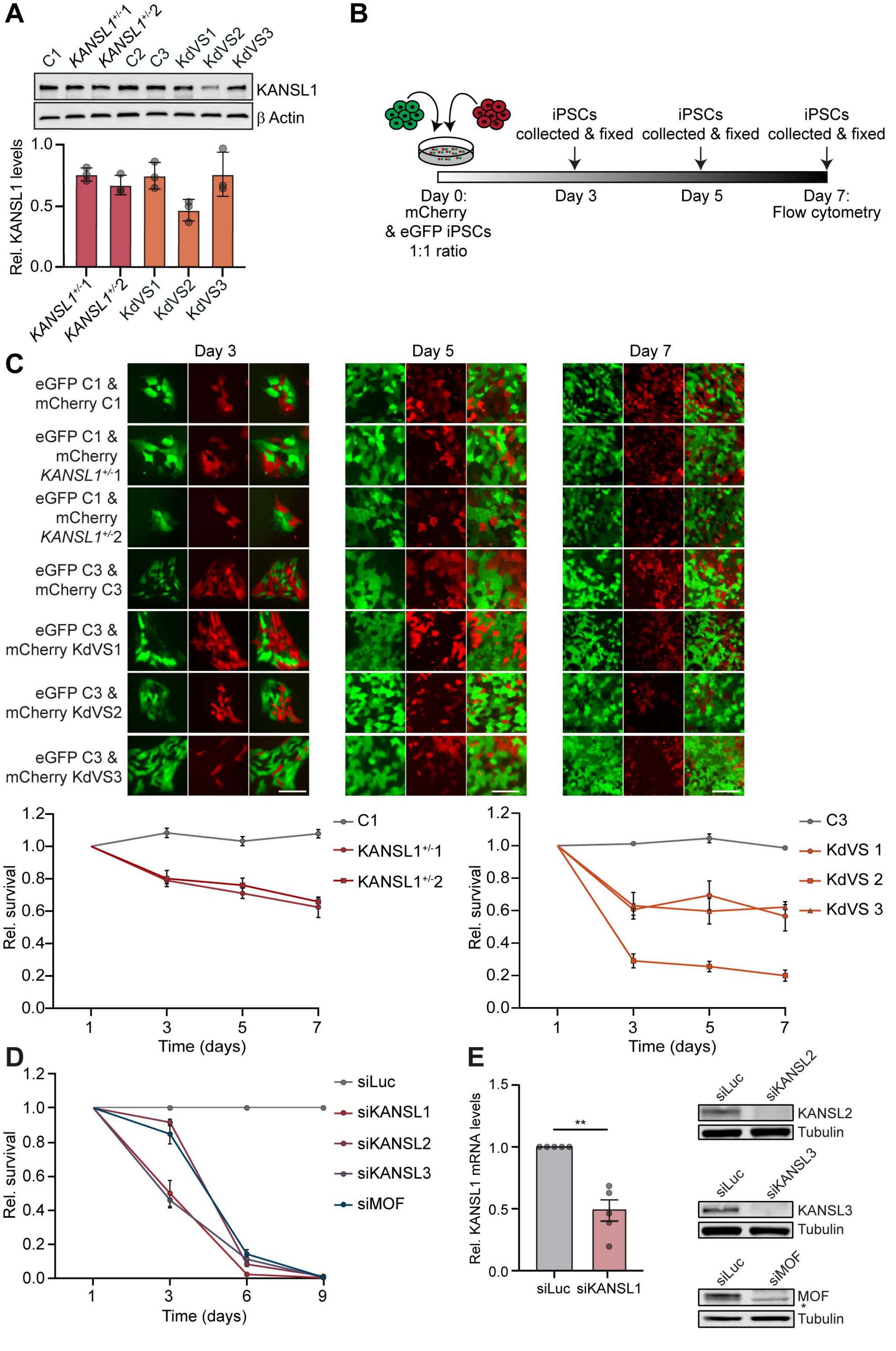

**Extended Data Fig. 1.**

A Western blotting analysis of KANSL1 in all iPSC lines (upper panel). Quantification of KANSL1 (121 kDa) expression relative to the loading control protein β actin (42 kDa) is shown (lower panel). Black circles represent KANSL1 protein quantification per experiment. N=3 independent experiments; mean ±SD.

B Competition assay protocol and time line for iPSCs. Cells were plated on day 0 in a 1:1 ratio. Cell ratio was measured on day 3, 5 and 7 post plating with flow cytometry.

C Representative images (upper panel) and quantification of competition assay of eGFP+ control, and mCherry+ KANSL1$^{+/-}$ and KdVS iPSCs (lower panels). Proliferation of *KANSL1*$^{+/-}$ and KdVS iPSCs are shown normalized to C1 and C3, respectively, which was set to 1. N=3 independent experiments. Circles represent the mean ±SEM. Scale bars 100µm.

D Competition assay measuring the relative proliferation after depletion of the NSL complex in GFP + and dsRED + U2OS cells at different timepoints. Proliferation was normalized to siLuc, which was set to 1. N=3 independent experiments. Circles represent the mean ±SEM.

E Relative KANSL1 mRNA levels indicated by RT-qPCR in U2OS cells, 2 days after treatment with the indicated siRNAs (left). KANSL1 expression was normalised to siLuc, which was set to 1. N=5 independent experiments. Black circles represent the mean of independent experiments; mean ± SEM plotted; unpaired two-tailed t test (** p<0.01). Western blot indicating protein expression levels of KANSL2, KANSL3 and MOF in U2OS cells 6 days after treatment with the indicated siRNAs (right). One representative experiment of two experiments shown.

Lingeman et al_Extended Data Figure 2

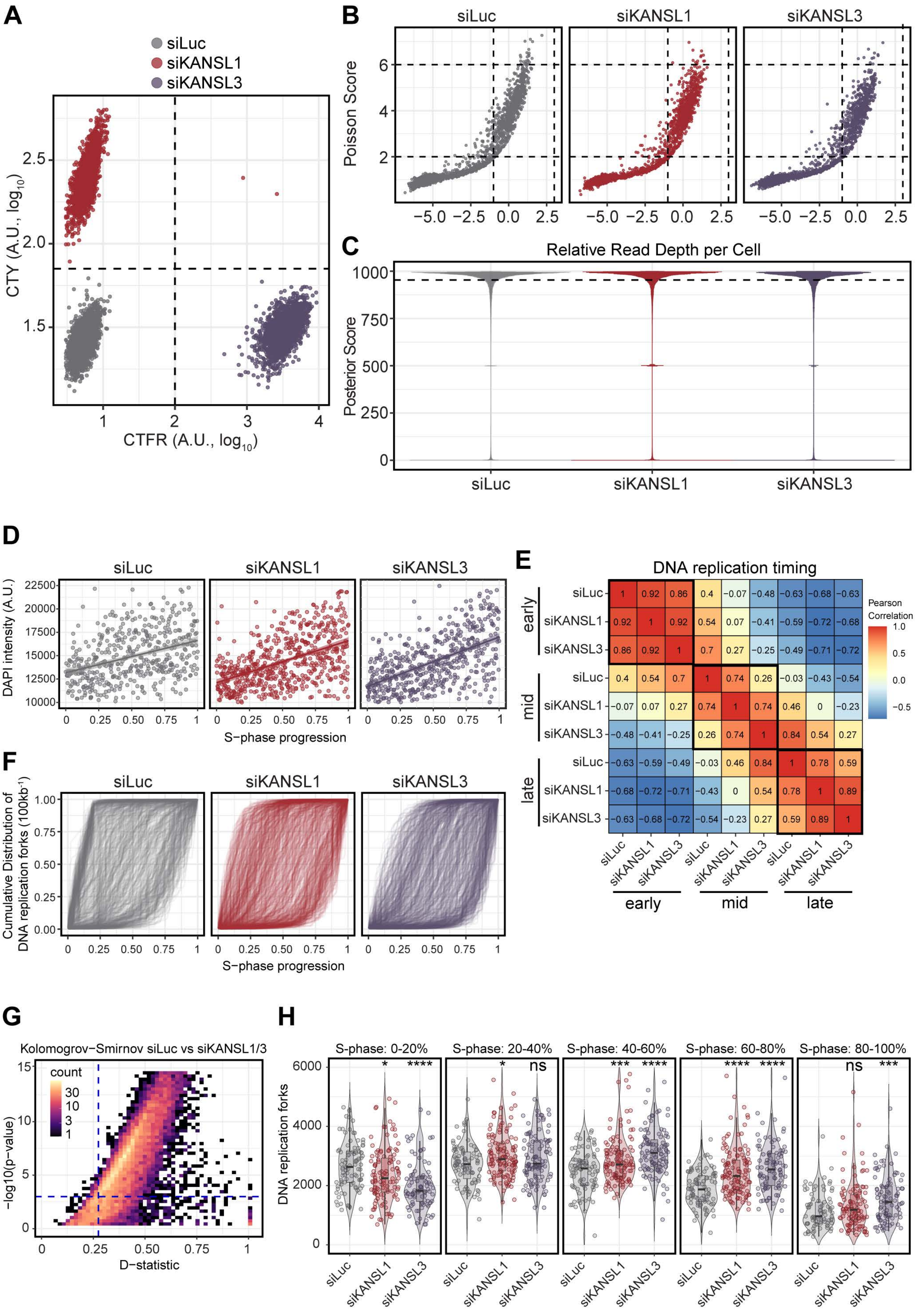

**Extended Data Fig. 2.**

A Flow cytometry data of U2OS cells 4 days after treatment with the indicated siRNAs for CellTrace labeled populations with mock (siLuc) CellTrace Yellow intensity (siKANSL1, y-axis) and CellTrace Far Red (siKANSL3, x-axis). Transfection conditions (colored circles) were determined by the gates indicated with the dashed lines. Cells in the upper right quadrant were excluded for any downstream analysis as we could not determine their population of origin.

B Poisson score (y-axis) versus average relative reads per bin (x-axis) for transfected cells. The right, middle area between four dashed lines contains cells selected for subsequent analysis. Each circle represents a single cell.

C Violin plot of posterior probability derived from HMM and Viterbi algorithm for U2OS cells transfected with the indicated siRNAs. All reads were considered for and subjected to scEdU-seq.

D Scatter plot of S-phase progression (x-axis) and DAPI intensity (y-axis) for U2OS cells transfected with the indicated siRNAs. The line indicates a linear fit with confidence interval.

E Pearson correlation matrix representation for each replication timing position (i.e., early, mid and late) for U2OS cells transfected with the indicated siRNAs. Number and colors indicate Pearson correlation

F The Cumulative frequency distribution (y-axis) of DNA replication forks for all 100kb bins on Chromosome 2 over S-phase progression (x-axis).

G Kolmogorov-Smirnov test for different Cumulative frequency distributions for siLuc versus siKANSL1 and siLuc versus siKANSL3 with the D-statistic (i.e., effect size, x-axis) and the -$\log_{10}$(adj.p-value) (i.e., significance, y-axis)

H The number of DNA replication forks (x-axis) for siLuc, siKANSL1 and siKANSL3 transfected U2OS (y-axis) cells split by S-phase Progression (bins for each 20%). The box of the boxplot is defined by the median ± IQR and whiskers are 1.5X IQR. Two-tailed t-test with multiple testing correction was performed (ns = not significan, * p<0.05, *** p<0.001, **** p<0.0001).

Lingeman et al_Extended Data Figure 3

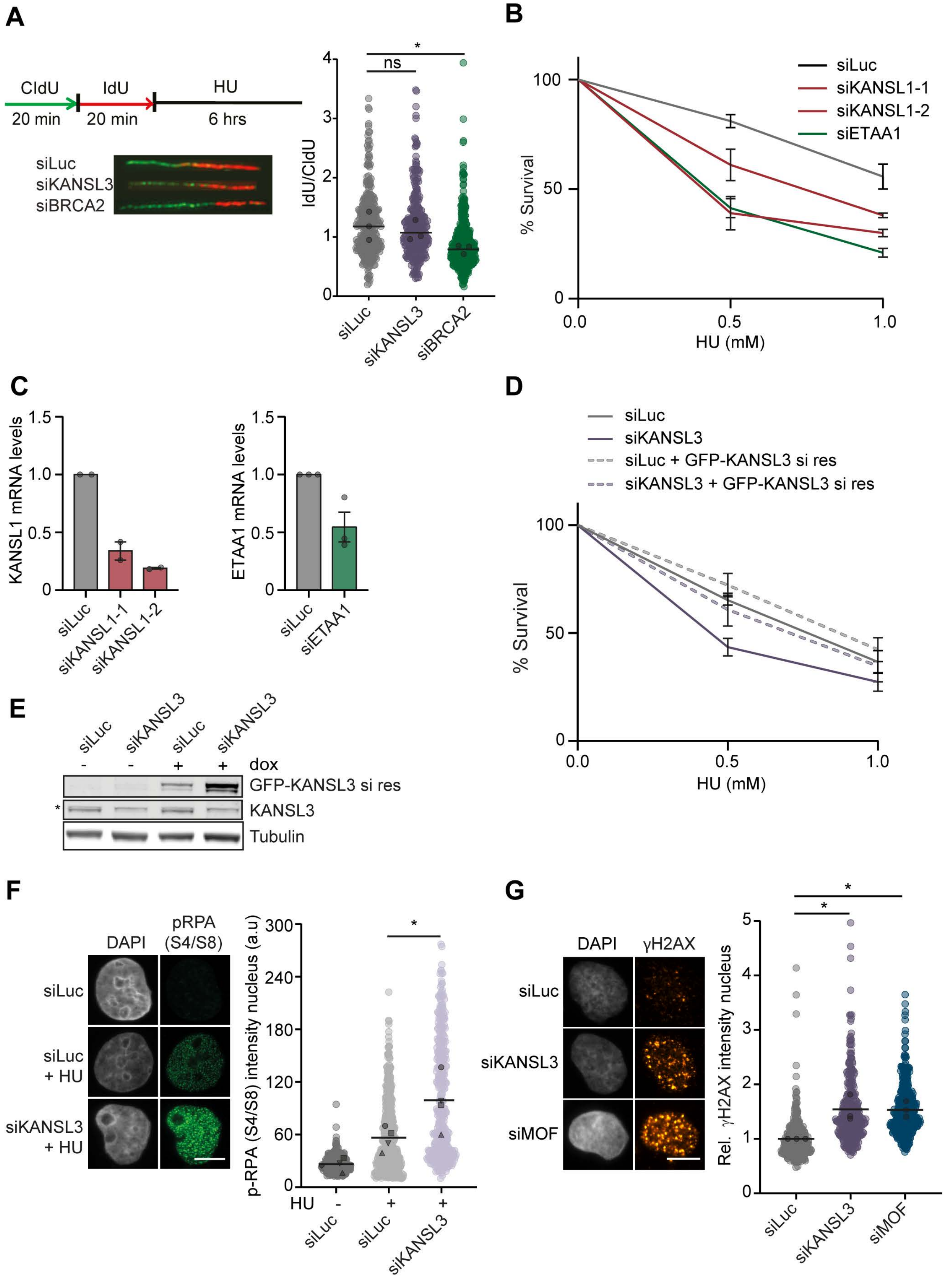

**Extended Data Fig. 3.**

A Labelling scheme, representative fibers (left) and quantification of DNA fork protection assay (right) in U2OS cells, 2 days after treatment with the indicated siRNAs. After Cldu and IdU labelling cells were treated with 4 mM HU for 6h. N=3 independent experiments. Black line represents the median of all data points, black circles the median of the independent experiments; one-way ANOVA and Dunnett's multiple comparisons test on median independent experiments (* $p<0.05$).

B Relative clonogenic survival in HCT116 cells after treatment with the indicated siRNAs, after treatment with varying doses of HU. Survival is relative to siLuc which was set to 100. N=3 independent experiments. Mean ±SEM indicated.

C Relative KANSL1 mRNA expression of cells described in B measured via RT-qPCR. N=2 independent experiments. Black circles represent the mean of independent experiments. Mean ±SEM indicated.

D Relative clonogenic survival in HCT116 cells after treatment with the indicated siRNAs, after treatment with varying doses of HU. Complementation of siRNA by doxycycline-induced overexpression of siRNA resistant (si res) GFP-KANSL3. N=2 independent experiments. Mean ±SEM indicated.

E KANSL3 expression levels of cells described in D, indicated by western blot. Endogenous KANSL3, GFP-KANSL3 siRNA resistant shown and Tubulin (loading control) shown. * marks an a-specific band.

F Representative images (right) and pRPA (S4/S8) levels (left) after HU treatment in KANSL3-depleted cells, treated for 4 hours +/- 2mM HU 2 days after siRNA treatment with the indicated siRNAs. N=6 independent experiments. Black line represents the mean of all data points, black circles the mean of the independent experiments; paired t-test on the means of independent experiments (* $p<0.05$, ** $p<0.01$). Scale bars 10 μm.

G Representative images (right) and relative γH2AX mean intensity levels (left) in U2OS cells, 2 days after treatment with the indicated siRNAs. γH2AX mean intensity levels were normalised to siLuc per experiment, which was set to 1. Black line represents the mean of all data points, black circles the mean of the independent experiments; one-way ANOVA and Dunnett's multiple comparisons test on mean independent experiments (* $p<0.05$). Scale bars 10 μm.

Lingeman et al_Extended Data Figure 4

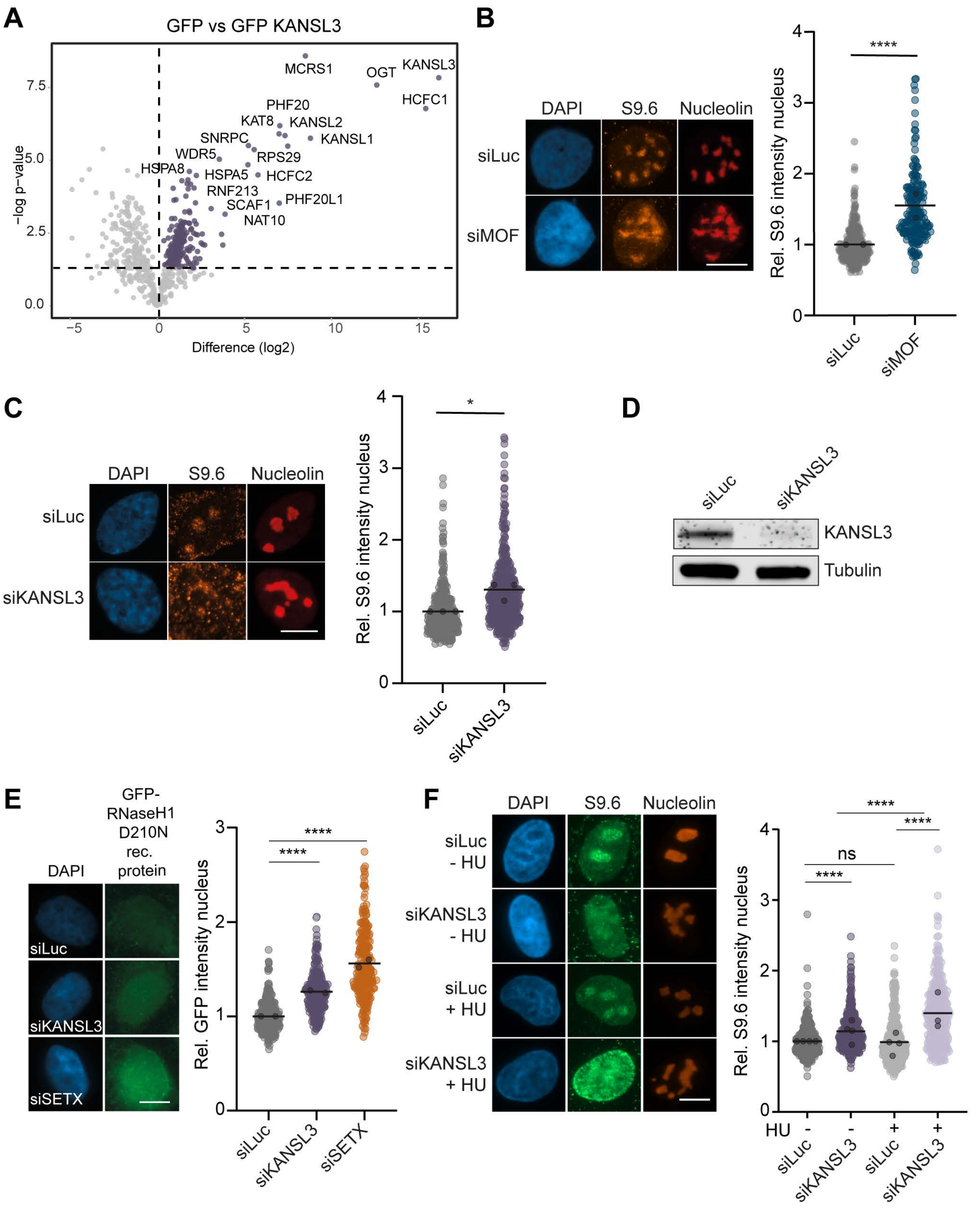

**Extended Data Fig. 4.**

A Mass spectrometry for GFP-KANSL3 vs GFP in U2OS cells. Purple circles represent proteins interacting with GFP-KANSL3. Top hits are annotated.

B Representative images (left) and relative R-loop analysis (right) in MOF-depleted U2OS cells, 6 days after treatment with the indicated siRNAs. Mean S9.6 intensity per nucleus (excluding the nucleolar signal) was normalized to siLuc, which was set to 1. N=2 independent experiments. Black line represents the mean of all data points, black circles the mean of the independent experiments; Kruskall Wallis test with Dunn's multiple comparisons test on all data points (**** p<0.0001). Scale bar 10 µm.

C Representative pictures (left) and relative R-loop analysis (right) in KANSL3-depleted RPE1 cells, 2 days after treatment with the indicated siRNAs. Mean S9.6 intensity per nucleus (excluding the nucleolar signal) was normalized to siLuc, which was set to 1. N=3 independent experiments. Black line represents the mean of all data points, black circles the mean of the independent experiments; unpaired two-tailed t test on means independent experiments. (* p<0.05). Scale bar 10 µm.

D Western blot confirming KANSL3 depletion in RPE1 cells, Tubulin was stained for as loading control. One representative experiment of two experiments shown.

E Representative images (left) and R-loop analysis (right) by GFP-RNaseH1 D210N protein staining in U2OS cells, 2 days after treatment with the indicated siRNAs. Mean GFP levels were normalized to siLuc, which was set to 1. N=2 independent experiments. Black line represents the mean of all data points, black circles the mean of the independent experiments; Kruskall Wallis test with Dunn's multiple comparisons test on all data points (**** p<0.0001). Scale bar 10 µm.

F Representative images (left) and R-loop analysis (right) by S9.6 antibody in U2OS cells treated for 4 hours +/- 2mM HU, 2 days after treatment with the indicated siRNAs. Mean S9.6 per nucleus excluding nucleolar signal was normalized to siLuc, which was set to 1. N=3-4 independent experiments. Black line represents the mean of all data points, black circles the means of the independent experiments; Kruskall Wallis test with Dunn's multiple comparisons test on all data points (ns = not significant, **** p<0.0001).

### Lingeman et al_Extended Data Figure 5

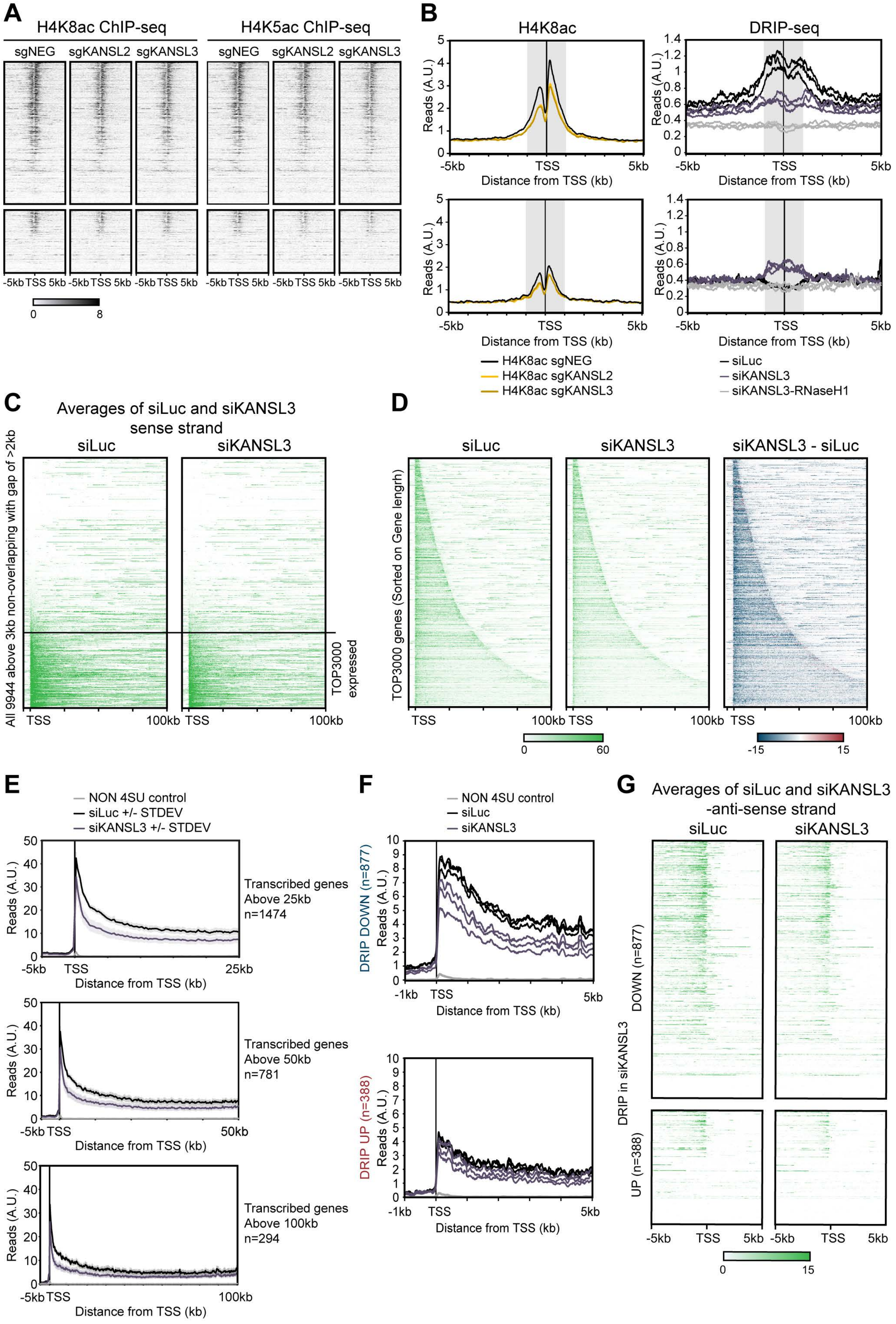

**Extended Data Fig. 5**

A Heatmaps of H4K8ac and H4K5ac ChIP-seq signals at indicated regions around TSSs of genes of at least 5kb that show decreased (DRIP-DOWN; n=877) or increased (DRIP-UP; n=388) DRIP signal at their TSS (+/-1 kb) upon KANSL3 depletion compared to controls. H4K5ac and K4K8ac ChIP-seq data previously published[1].

B Metaprofiles of H4K8ac ChIP-seq and DRIP-seq signals around TSSs with decreased (DOWN) or increased (UP) DRIP signal in siKANSL3 compared to siLuc. The grey shaded areas are regions from -1kb to +1kb around the TSS used to quantify decreased or increased DRIP signal.

C Heatmaps of TTchem-seq signals in cells treated with siLuc or siKANSL3 at indicated regions around 9944 TSSs of genes of at least 3kb with at least 2kb between genes. Data presents signal of sense transcription and heatmaps are sorted on TTchem-seq signal in the first 3kb of genes. Indicated are the TOP3000 TSSs defined as being actively transcribed.

D Heatmaps of TTchem-seq signals in cells treated with siLuc or siKANSL3 at indicated regions around the TOP3000 actively transcribed TSSs defined in C. Data presents signal of sense transcription and heatmaps are sorted on gene length.

E Metaprofiles of TTchem-seq signals around actively transcribed TSSs of genes of at least 25 kb (top) 50 kb (middle) or 100 kb (bottom).

F Metaprofiles of DRIP-seq signals around TSSs, as separate replicates, with decreased (DOWN) or increased (UP) DRIP signal in siKANSL3 compared to siLuc as in B.

G Heatmaps of antisense TTchem-seq signals at indicated regions around TSSs that show decreased (DRIP-DOWN; n=877) or increased (DRIP-UP; n=388) DRIP signal upon KANSL3 depletion compared to controls.

Lingeman et al_Extended Data Figure 6

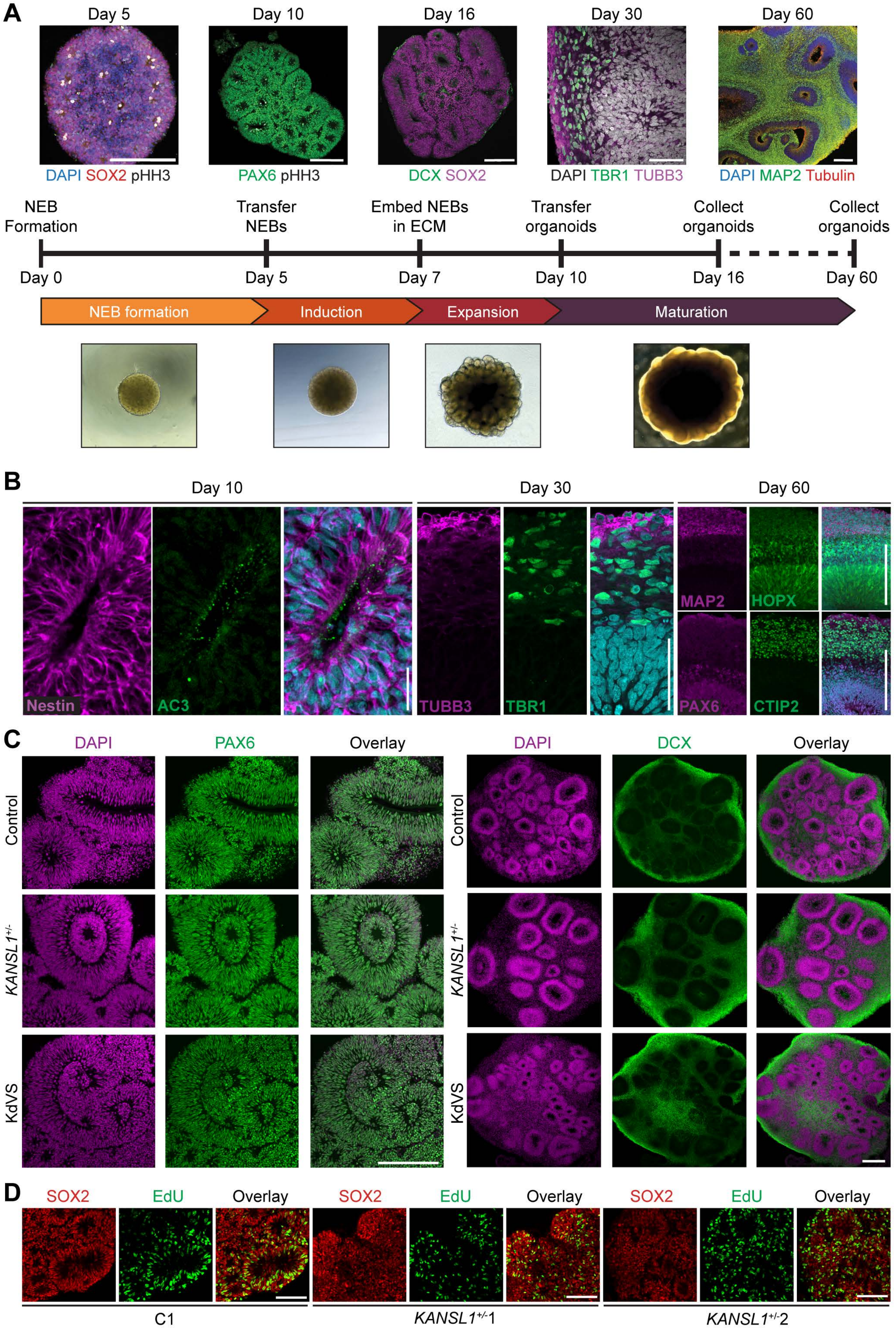

**Extended Data Fig. 6.**

A Neural organoids were differentiated from iPSCs via neural epithelial bodies (NEBs) into neural organoids over the course of 60 days and assessed at day 16 and day 60. Immunofluorescent images of control organoids over development are shown above the protocol schematic and bright field images of the organoids are shown below. The NEB on day 5 (formation) image is stained with the mitotic marker phosphorylated histone H3 (white), the radial glia / neuroprogenitor marker SOX2 (red), DAPI (blue). Day 10 (expansion) shows the formation of polarized neuroepithelial rosettes stained with the radial glia marker PAX6 (green). The day 16 organoid is stained with SOX2 (magenta) and the migrating neuroblast marker DCX (green). Day 30 shows an organoid stained with the general neuronal marker TUBB3 (magenta) and neuronal marker TBR1, scale bars are 50 µm. Day 60 (maturation) shows an overlay of the microtubule marker tubulin (red) and dendritic marker MAP2 (green). Scale bars for all images are 200 µm unless otherwise noted.

B All neural organoid rounds were validated for expression of neurodevelopmental markers and appropriate tissue architecture on day 10, 30, and 60 of differentiation. Day 10, Scale bars are 25 µm . Day 30, 60 scale bar are 50 µm.

C Control, *KANSL1*$^{+/-}$, and KdVS organoids differentiated along similar trajectories based on architecture and expression of neural markers. Scale bars are 100 µm.

D Wide field image of EdU integration in C1 and *KANSL1*$^{+/-}$ organoids on day 16. Scale bars are 100 µm.

### Lingeman et al_Extended Data Figure 7

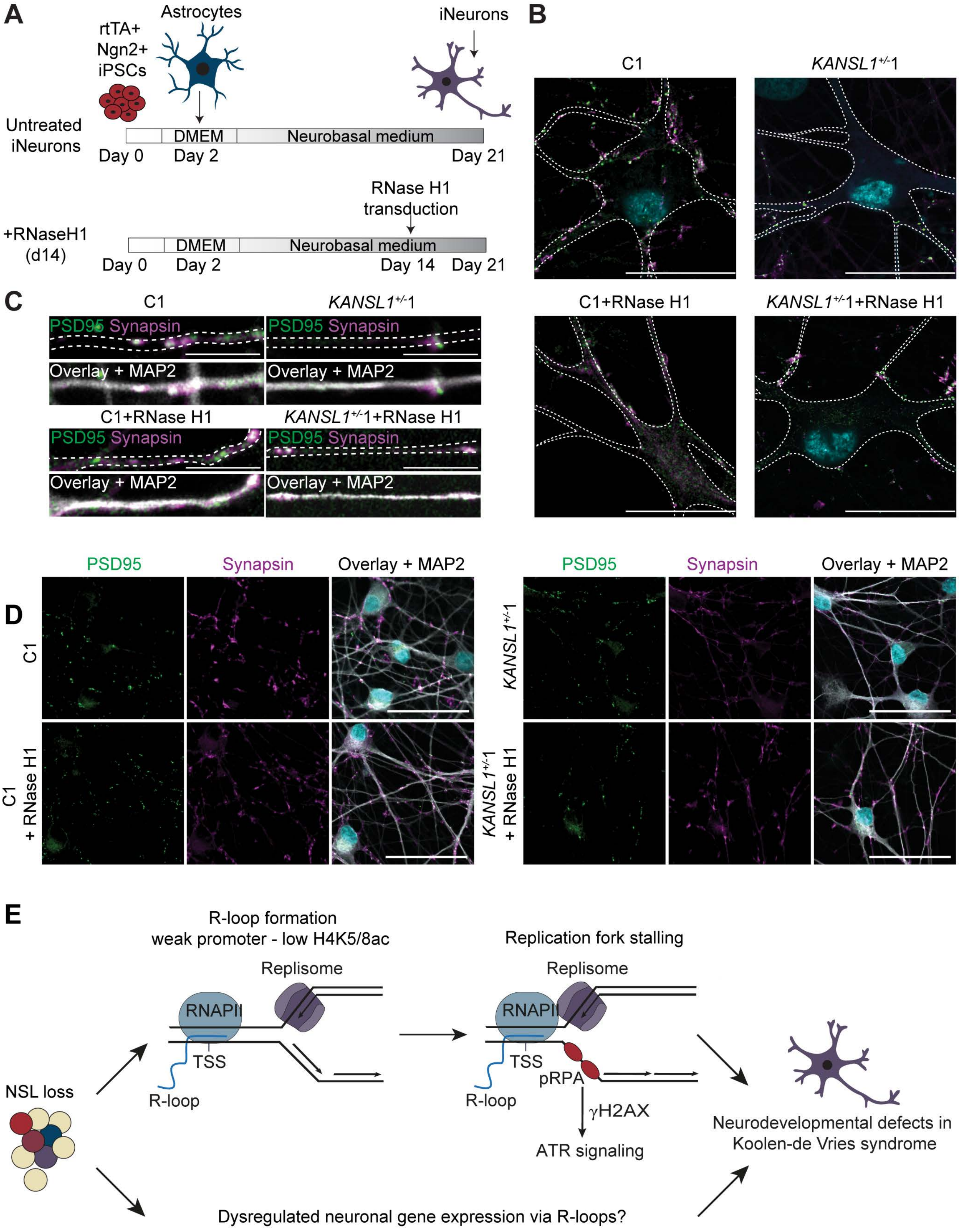

**Extended Data Fig.7.**

A Schematic diagram of iNeuron differentiation protocol in untreated neurons and neurons transduced with RNase H1 on day 14 of differentiation.

B Representative image of iNeuron traces to show localization of functional synapses. The presynaptic marker Synapsin I (magenta) and the post synaptic marker PSD95 (green) are shown along the dendrites and soma of the outlined neuron. Outline was drawn based on MAP2 staining. Scale bars are 25 µm.

C Overlay of the presynaptic marker Synapsin I (magenta) and the post synaptic marker PSD95 (green), and the dendritic marker MAP2 (white) on of dendrites of iNeurons differentiated for 21 days. Traces of the dendrite are shown as white outlines for visualization of the pre- and post-synaptic colocalization, in the overlay MAP2 is included to show the structure of the dendrite. Scale bars are 10 µm.

D Representative wide view images of C1 and *KANSL1*$^{+/-}$1 day 21 iNeurons +/- RNase H1 treatment on day 14 during differentiation. PSD95 is shown in green, Synapsin in magenta, and the overlay additional illustrates MAP2 in white and DAPI in cyan. Scale bar 50 µm.

E Model of how altered DNA-RNA hybrid levels influence neuronal differentiation and maturation in KdVS.

**References**

1. Radzisheuskaya, A. *et al.* Complex-dependent histone acetyltransferase activity of KAT8 determines its role in transcription and cellular homeostasis. *Mol Cell* **81**, 1749-1765.e1748 (2021).

**Supplementary Table 1. List of oligos**

| siRNAs / gRNAs | Sequence |
|---|---|
| siLuc | CGUACGCGGAAUACUUCGA |
| siKANSL1-1 | CGGCAACGCCAACAUCCUU |
| siKANSL1(-2) | GAAGCGGAGGCUUGUUCGA |
| siKANSL2 | CAGUGAAGCCAGCCGAAUA |
| siKANSL3 | UGAUGACAAUCUCAGAAUA |
| siMOF | CAAGAUCACUCGCAACCAA |
| siRAD51 | GAGCUUGACAAACUACUUC |
| siSETX | GCCAGAUCGUAUACAAUUAUU |
| siBRCA2 | GAAGAAUGCAGGUUUAAUA |
| siETAA1 | GAGCAAAACAAGAGGAAUUUU |
| *KANSL1*$^{+/-}$1 target sequence (gRNA) | CTTAGAACCATGAATACGAG |
| *KANSL1*$^{+/-}$2 target sequence (gRNA) | GAGCCCGTTTTCCCCCATTG |

**Supplementary Table 2. List of antibodies**

| Antibody | Source | Identifier |
|---|---|---|
| Adenylate Cyclase 3 (ADCY3) (host: rabbit, used 1:500 IF[1]) | ProteinTech | Cat#19492-1-AP, RRID AB_10638445 |
| Alexa Fluor 647 anti-SOX2 (host: mouse, used 1/200 FC[2]) | BioLegend | Cat#656108, RRID AB_2563681 |
| B-actin (host: mouse, used 1/5000 WB[3]) | Invitrogen | Cat#MA1-140, RRID AB_2536844 |
| CldU / Anti-BrdU, BU1/75 (host: rat, used 1/500 IF) | Abcam | Cat#Ab6326, RRID AB_2313786 |
| CTIP2 (host: rat, used 1/200 IF) | Abcam | Cat#Ab18465, RRID AB_2064130 |
| Doublecortin (host: mouse, used 1/500 IF) | Santa Cruz Biotechnologies | Cat#sc-271390, RRID AB_10610966 |
| FluoTag-X2 anti-PSD95 (host: alpaca, used 1/200 IF) | NanoTag Biotechnology | Cat#N3702-AF647-L, RRID AB_2936216 |
| GFP (host: Mouse, used 1/2500 WB) | Sigma-Aldrich | Cat#11814460001, RRID AB_390913 |
| HOPX (host: rabbit, used 1/200 IF) | ProteinTech | Cat#11419-1-AP, RRID AB_10693525 |
| IdU / Anti-BrdU, B44 (host: mouse, used 1/700 IF) | Bio-Rad | Cat#Ab6326, RRID AB_2313786 |
| KANSL1 (host: rabbit, used 1/500 WB) | Sigma-Aldrich | Cat#HPA006874, RRID:AB_1852393 |
| KANSL2 (host: rabbit, used 1/1000 WB) | ATLAS | Cat#HPA038497, RRID AB_10674685 |
| KANSL3 (host: rabbit, used 1/2000 WB) | Sigma-Aldrich | Cat#HPA035018, RRID AB_10601763 |
| MAP2 (host: guinea pig, used 1/1000 IF) | Synaptic Systems | Cat#188 004, RRID AB_2138181 |
| MOF (host: rabbit, used 1/1000 WB) | Bethyl | Cat#A300-992A, RRID AB_805802 |
| Nestin (host: mouse, used 1/300 IF) | Invitrogen | Cat#MA1-110, RRID AB_2536821 |
| Nucleolin (host: rabbit, used 1/5000 IF) | Abcam | Cat#ab50279, RRID AB_881762 |
| PAX6 (host: rabbit, used 1/200 IF) | BioLegend | Cat#901301, RRID AB_2565003 |
| p-HH3 (S10) (host: mouse, used 1/50 IF) | Invitrogen | Cat#MA515220, RRID AB_11008586 |
| p-RPA (S33) (host: rabbit, used 1:1000 IF) | Bethyl | Cat#A300-246A, RRID AB_2180847 |
| p-RPA32 (S4/S8) (host: rabbit, used 1:1000 IF) | Bethyl | Cat#A300-245A, RRID AB_210547 |
| RNaseH1 (host: rabbit, used 1:1000 WB) | ProteinTech | Cat#15606-I-AP, RRID AB_2238624 |
| S9.6 (host: mouse, used 1:1000 IF) | Millipore | Cat#MABE1095, RRID AB_2861387 |
| SATB2 (host: mouse, used 1/500 IF) | Abcam | Cat#Ab51502, RRID AB_882455 |
| SOX2 (host: rabbit, used 1/200 IF) | Abcam | Cat#Ab97959, RRID AB_2341193 |

| | | |
|---|---|---|
| Synapsin I (host: rabbit, used 1/500 IF) | Sigma-Aldrich | Cat#AB1543P, RRID AB_90757 |
| TBR1 (host: rabbit, used 1/200 IF) | Abcam | Cat#ab183032, RRID AB_2936859 |
| TBR2 / Eomes (host: rabbit, used 1/200 IF) | Abcam | Cat#Ab23345, RRID AB_778267 |
| TUBB3 (host: mouse, used 1/500 IF) | BioLegend | Cat#801201, RRID AB_2313773 |
| Tubulin, Clone DM1A (host: mouse, used 1:1000 WB) | Sigma-Aldrich | Cat#T6199, RRID AB_477583 |
| γH2AX (S139) (host: mouse, used 1:1000 IF) | Millipore | Cat#05-636, RRID AB_309864 |
| goat anti-mouse Ig (host: goat, used 1/1000 WB) | Biotium | Cat#CF770, RRID AB_10559194 |
| goat anti-rabbit Ig (host: goat, used 1/1000 WB) | Biotium | Cat#CF680,RRID AB_10557108 |
| HRP goat anti-mouse (host: goat, used 1/5000 WB) | Jackons ImmunoResearch | Cat#115-035-062, RRID AB_2338504 |
| HRP goat anti-mouse (host: goat, used 1/5000 WB) | Invitrogen | Cat#G21234, RRID AB_2536530 |
| Goat anti-Guinea pig IgG (H+L) Highly Cross-Adsorbed Secondary Antibody, Alexa Fluor 488 (host: goat, used 1/500 IF) | Invitrogen | Cat#A-11073, RRID AB_2534117 |
| Goat anti-Mouse IgG (H+L) Highly Cross-Adsorbed Secondary Antibody, Alexa Fluor 488 (host: goat, used 1/500 IF) | Invitrogen | Cat#A-11029, RRID AB_2534088 |
| Goat anti-Rabbit IgG (H+L) Cross-Adsorbed Secondary Antibody, Alexa Fluor 488 (host: goat, used 1/500 IF) | Invitrogen | Cat#A-11008, RRID AB_143165 |
| Goat anti-Rat IgG (H+L) Cross-Adsorbed Secondary Antibody, Alexa Fluor 488 (host: goat, used 1/500 IF) | Invitrogen | Cat#A-11006, RRID AB_2534074 |
| Goat anti-Mouse IgG (H+L) Highly Cross-Adsorbed Secondary Antibody, Alexa Fluor 568 (host: goat, used 1/500 IF) | Invitrogen | Cat#A-11031, RRID AB_144696 |
| Goat anti-Rabbit IgG (H+L) Cross-Adsorbed Secondary Antibody, Alexa Fluor 568 (host: goat, used 1/500 IF) | Invitrogen | Cat#A-11011, RRID AB_143157 |
| Goat anti-Guinea pig IgG (H+L) Highly Cross-Adsorbed Secondary Antibody, Alexa Fluor 647 (host: goat, used 1/500 IF) | Invitrogen | Cat#A-21450, RRID AB_141882 |
| Goat anti-Mouse IgG (H+L) Cross-Adsorbed Secondary Antibody, Alexa Fluor 647 (host: goat, used 1/500 IF) | Invitrogen | Cat#A-21237, RRID AB_1500743 |
| Goat anti-Rabbit IgG (H+L) Highly Cross-Adsorbed Secondary Antibody, Alexa Fluor 647 (host: goat, used 1/500 IF) | Invitrogen | Cat#A-21245, RRID AB_2535813 |

[1] IF = immunofluorescence, [2] FC = flow cytometry, [3] WB = western blot

Lingeman et al_Supplementary Figure 1

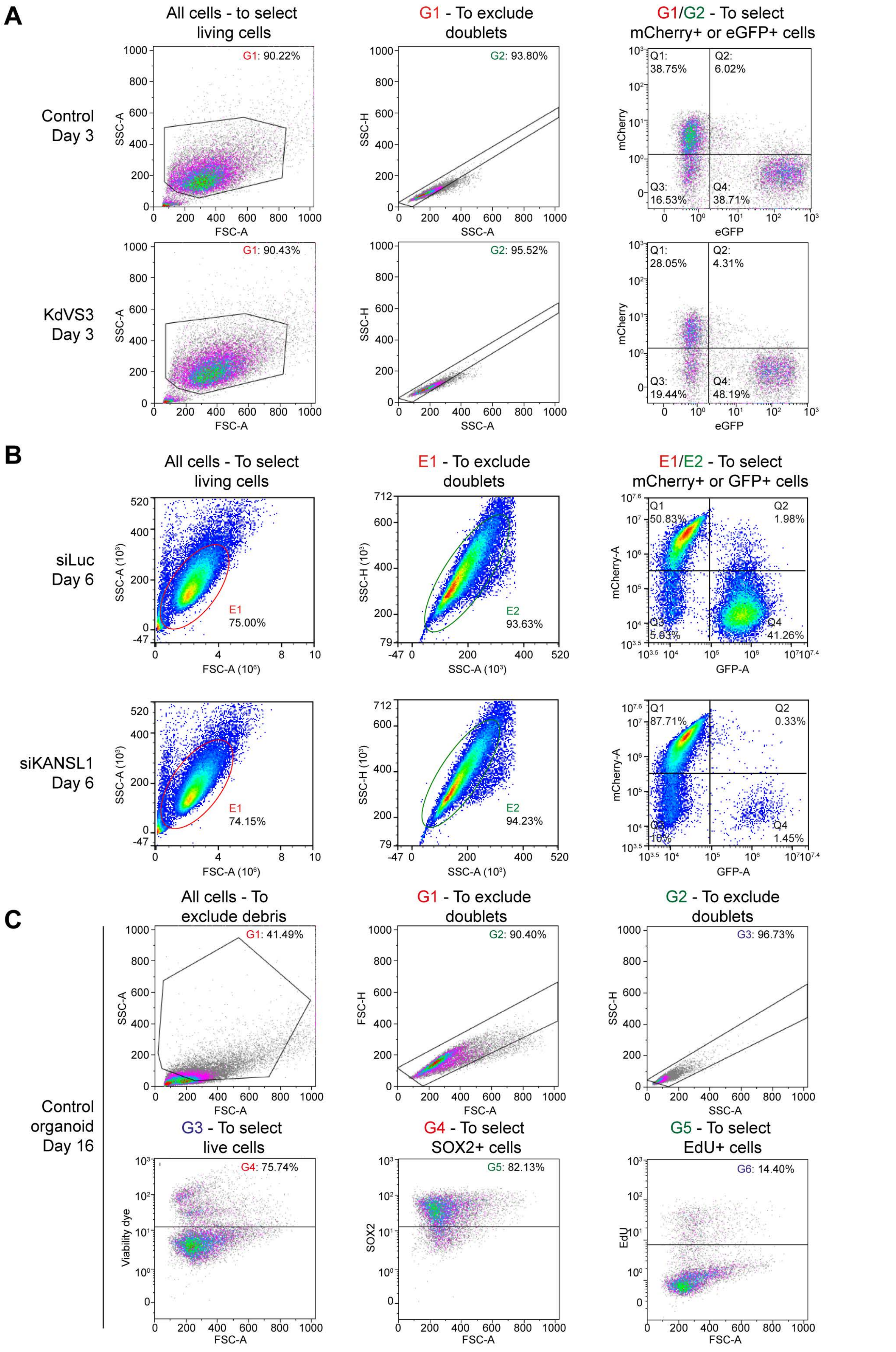

**Supplementary Fig. 1.**

A Representative flow cytometry dot plots showing the gating strategy for the growth competition assay with iPSCs derived from KdVS patients in Extended Data Fig. 1C.

B Representative flow cytometry dot plots showing the gating strategy for the growth competition assay with U2OS cells in Extended Data Fig. 1D.

C Representative flow cytometry dot plots showing the gating strategy for the EdU incorporation experiments with organoids shown in Fig. 7D.
